# Evaluating Extraction Methods to Study Canine Urine Microbiota

**DOI:** 10.1101/2021.01.15.425942

**Authors:** R. Mrofchak, C. Madden, M.V. Evans, V.L. Hale

**Author notes:** Corresponding Author: Vanessa Hale, Department of Veterinary Preventive Medicine, Ohio State University College of Veterinary Medicine, 1900 Coffey Rd., Columbus, OH 43210.

## Abstract

The urinary microbiota is the collection of microbes present in urine that play a role in host health. Studies of urine microbiota have traditionally relied upon culturing methods aimed at identifying pathogens. However, recent culture-free sequencing studies of the urine microbiota have determined that a diverse array of microbes are present in health and disease. To study these microbes and their potential role in diseases like bladder cancer or interstitial cystitis, consistent extraction and detection of microbial DNA from urine is critical. However, urine is a low biomass substrate, requiring sensitive methods to capture DNA and making the risk of contamination high. To address this challenge, we collected urine samples from ten healthy dogs and extracted DNA from each sample using five different commercially available extraction methods. Extraction methods were compared based on total and bacterial DNA concentrations and microbial community composition and diversity assessed through 16S rRNA gene sequencing. Significant differences in the urinary microbiota were observed by dog and sex but not extraction method. The Bacteremia kit yielded the highest total DNA concentrations (Kruskal-Wallis, *p* = 0.165, not significant) and the highest bacterial DNA concentrations (Kruskal-Wallis, *p* = 0.044). Bacteremia also extracted bacterial DNA from the greatest number of samples. Taken together, these results suggest that the Bacteremia kit is an effective option for studying the urine microbiota. This work lays the foundation to study the urine microbiome in a wide range of urogenital diseases in dogs and other species.

**Highlights:**

- Canine urine microbiota differed by sex and dog but not extraction method.
- Qiagen Bacteremia kit yielded the highest bacterial DNA concentrations from urine.
- The Bacteremia kit extracted bacterial DNA from the greatest number of samples.
- Absolute abundance of *Sphingomonas* species increased in female dog urine.
- *Pasteurellaceae* bacterium canine oral taxon 272 increased in male dog urine.

## Introduction

Urine, in the absence of urinary tract infection, has long been considered sterile; a principle still taught in many healthcare professional settings to date. However, evidence counter to this idea has been accumulating for several decades (K. Thomas-White et al., 2016). Culture-positive asymptomatic bacteriuria is commonly reported in women and older adults; although, this is sometimes deemed “contamination” based on bacterial counts < 10^5^ (Kass, 1962). Culture-negative symptomatic urinary tract infections (UTIs) are also common, and can, in some cases, be attributed to fastidious organisms that fail to grow using standard urine culturing (SUC) techniques (Dune et al., 2017; Kass, 1962; Maskell et al., 1979; Maskell, 2010; Price et al., 2016; Thapaliya et al., 2020; Thomas-White et al., 2018). More recently, culture-independent next-generation sequencing of urine and enhanced quantitative urine culture (EQUC) has revealed the presence of bacteria in >90% of individuals – including those with and without UTIs and from urine collected via free-catch, transurethral catheter, or suprapubic aspirates (Chen et al., 2018; Hilt et al., 2014; Pearce et al., 2014; Pohl et al., 2020; Price et al., 2020, 2016; Thapaliya et al., 2020; Thomas-White et al., 2018; Wolfe et al., 2012). Collectively, this work provides evidence for the presence of a urinary microbiome containing live bacteria that are present in healthy individuals and distinct from contaminants.

A growing body of work has revealed profound links between the microbiota (oral, gut, lung, vaginal) and host health (Erb-Downward et al., 2011; Fettweis et al., 2019; Griffen et al., 2012; Manor et al., 2020; Ravel et al., 2011) – from nutrient metabolism (David et al., 2014), to immune development and defense (Furusawa et al., 2013), to colonization resistance (Zmora et al., 2018). Thus, it follows that the urine / bladder microbiome may also have a critical role in host health, but it has been far less studied than the microbiota of the aforementioned body regions. Notably, the “urine is sterile” dogma contributed to the exclusion of urine in the first phase of Human Microbiome Project (HMP) launched in 2007 (Proctor et al., 2019). In 2014, the second phase of the HMP launched and included the urine / bladder microbiome. However, work on the urine microbiome continues to lag. Although urine is not sterile, it has a low microbial biomass, making it more challenging to characterize and at greater risk for contamination (Eisenhofer et al., 2019). Despite this, several recent studies on urine have identified clear shifts in the microbial community associated with age (Adebayo et al., 2020; Curtiss et al., 2018; Gottschick et al., 2017; Komesu et al., 2018; Liu et al., 2017b; Pearce et al., 2014), sex (Lewis et al., 2013; Pederzoli et al., 2020; Pohl et al., 2020), urgency urinary incontinence in women (Brubaker et al., 2014; Karstens et al., 2016; Pearce et al., 2014; K. J. Thomas-White et al., 2016), bladder cancer (Bi et al., 2019; Bučević Popović et al., 2018; Chipollini et al., 2020; Hourigan et al., 2020; Liu et al., 2019; Mansour et al., 2020; Pederzoli et al., 2020; Wu et al., 2018), interstitial cystitis (Abernethy et al., 2017; Bresler et al., 2019; Domingue et al., 1995; Meriwether et al., 2019; Nickel et al., 2019, 2016; Siddiqui et al., 2012), neuropathic bladders (Forster et al., 2020; Fouts et al., 2012) and pneumonia (Pierre et al., 2020). However, additional work is needed to identify definitive and mechanistic links between the urine / bladder microbiome, host health, and urinary tract disease.

Urinary tract disease is one of the most common diagnoses in veterinary medicine (Byron, 2019; Ling, 1984). In addition, dogs are a valuable translational model for many human diseases, including urogenital diseases like bladder cancer (Knapp et al., 2014). However, there has only been one study, to our knowledge, characterizing canine urine microbiota (Burton et al., 2017) and none evaluating canine urine DNA extraction methods. Multiple methods of urine microbial DNA isolation have been reported in human studies (Bergallo et al., 2006; El Bali et al., 2014; Fouts et al., 2012; Karstens et al., 2020; Pohl et al., 2020; Siddiqui et al., 2009), and there are a few studies that have compared microbial DNA extraction methods. These studies include a comparison of methods for extracting fungal (Ackerman et al., 2019) and viral DNA from urine (Bergallo et al., 2006; Buffone et al., 1991; Santiago-Rodriguez et al., 2015) as well as a study on methods to reverse crystal precipitates that interfere with DNA extraction (Munch et al., 2019). Yet, only one recent study has evaluated urine DNA extraction methods in relation to the bacterial microbiota in humans (Karstens et al., 2020), and there are no comparable studies in dogs, which are a valuable and translational model for human urinary tract diseases (Knapp et al., 2014). In this study, our objective was to compare canine urine bacterial DNA quantity and 16S rRNA sequencing results of five commonly used extraction methods. These extraction methods included four DNA isolation kits from Qiagen—Bacteremia, Blood and Tissue, PowerFecal, and PowerFecal Pro—and an extraction protocol using magnetic beads.

## Materials and Methods

### Sample collection

Mid-stream free catch urine samples were collected from 10 healthy dogs in September 2019 at The Ohio State University College of Veterinary Medicine (Columbus, OH, USA) with owner consent (IACUC #2019A00000005). Owners were given a questionnaire to provide information including age, sex, breed, spay / neuter status, and diet. Enrollment criteria included: Dogs had to weigh at least 30 lbs (∼13.6 kg) and produce > 30 ml of urine in a single urination. Dogs were excluded if they had any history of urinary tract disease or antibiotic use within three weeks of sample collection. We collected urine samples from a total of 10 dogs, including 4 males (3 neutered) and 6 females (5 spayed). The average age of the dogs was 3.7 years old (range: 0.75 – 10 years) and represented multiple breeds including: one Great Pyrenees, one Labrador Retriever, one Golden Retriever, and seven mixed breed dogs. Full metadata on each dog is available in **Table S1**. All urine samples were stored at −80 °C within 6 hours of urination where they remained until extraction.

### DNA extraction

Five extraction methods were tested: QIAamp^®^ BiOstic^®^ Bacteremia DNA Kit (B), DNeasy^®^ Blood and Tissue Kit (BTL), QIAamp^®^ PowerFecal^®^ DNA Kit (PF), QIAamp^®^ PowerFecal^®^ Pro DNA Kit (PFP) (Qiagen, Venlo, Netherlands) and an extraction protocol using magnetic beads (MB) (Liu et al., 2017a). Each protocol incorporated varying chemical, mechanical, and thermal lysing steps to facilitate DNA extraction (**Table 1**).

**Table 1:**
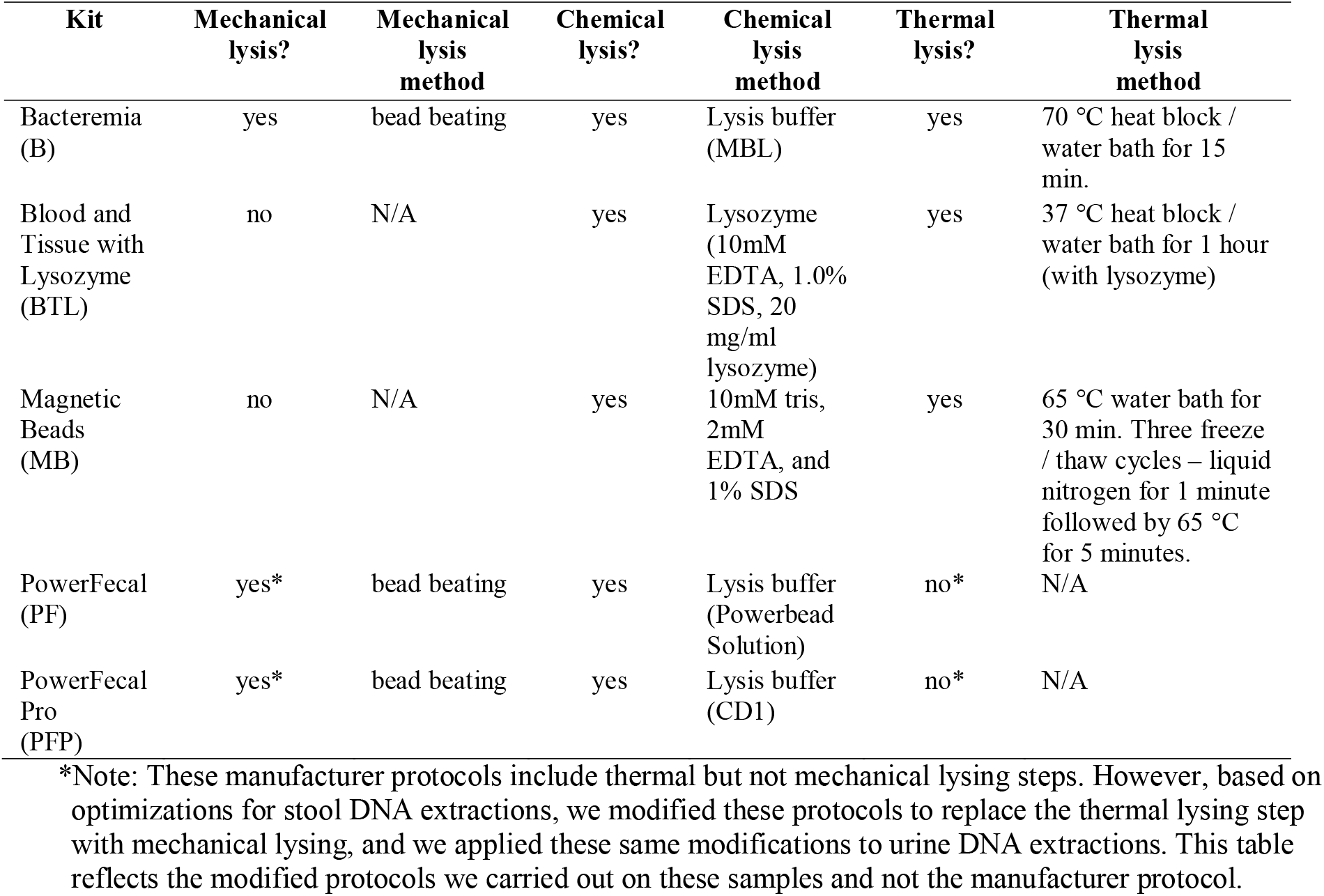
Mechanical, chemical and thermal lysis properties of each extraction method.

| Kit | Mechanical lysis? | Mechanical lysis method | Chemical lysis? | Chemical lysis method | Thermal lysis? | Thermal lysis method |
| --- | --- | --- | --- | --- | --- | --- |
| Bacteremia (B) | yes | bead beating | yes | Lysis buffer (MBL) | yes | 70 °C heat block / water bath for 15 min. |
| Blood and Tissue with Lysozyme (BTL) | no | N/A | yes | Lysozyme (10mM EDTA, 1.0% SDS, 20 mg/ml lysozyme) | yes | 37 °C heat block / water bath for 1 hour (with lysozyme) |
| Magnetic Beads (MB) | no | N/A | yes | 10mM tris, 2mM EDTA, and 1% SDS | yes | 65 °C water bath for 30 min. Three freeze / thaw cycles – liquid nitrogen for 1 minute followed by 65 °C for 5 minutes. |
| PowerFecal (PF) | yes* | bead beating | yes | Lysis buffer (Powerbead Solution) | no* | N/A |
| PowerFecal Pro (PFP) | yes* | bead beating | yes | Lysis buffer (CD1) | no* | N/A |

\*Note: These manufacturer protocols include thermal but not mechanical lysing steps. However, based on optimizations for stool DNA extractions, we modified these protocols to replace the thermal lysing step with mechanical lysing, and we applied these same modifications to urine DNA extractions. This table reflects the modified protocols we carried out on these samples and not the manufacturer protocol.

To prepare urine samples for extraction, 3.0 ml of urine was centrifuged at 4 °C and 20,000 x *g* for 30 minutes. Samples that underwent MB extraction were centrifuged at 4 °C and 20,000 x *g* for 20 minutes. After centrifuging, the supernatant was removed and discarded and the pellet was used for extraction. Samples were assigned initials unique to each dog (AW, CB, CS, DD, DH, HB, LS, SF, SM, and ZR) followed by the abbreviation of the extraction method (B, BTL, MB, PF, or PFP). For example, sample AWB is urine from dog AW extracted using the Bacteremia (B) kit. Negative and positive controls were also extracted from each method. The negative control was a blank (no sample) tube that underwent the full extraction process. The positive control was 3.0 ml of urine from dog AW spiked with 3 × 10^8^ CFUs of *Melissococcus plutonius. M. plutonius* is a honeybee pathogen that would not be expected in the urine / gut of dogs (Djukic et al., 2018). Brief descriptions of each extraction method are included below.

### Bacteremia

Urine pellets were resuspended in a lysis buffer (MBL) and placed in a 70 °C water bath for 15 minutes. Samples then underwent two rounds of bead beating (6 m/s for 60 seconds with a 5 minute rest between rounds). Bead beating was performed on a MP FastPrep-24™ 5G (MP Biomedicals, Santa Ana, California, USA). After bead beating, the samples were cleaned with an Inhibitor Removal Solution. The remainder of the protocol was followed with two modifications. First, centrifugation was performed at 13,000 x *g* instead of 10,000 x *g*. Second, during the final step, DNA was eluted into 50 μl of elution buffer (EB) and incubated at room temperature for five minutes; then, the eluent was run through the silica membrane of the spin column a second time to maximize DNA yield.

### Blood and Tissue with Lysozyme

Urine pellets were resuspended in a lysis buffer adapted from Pearce *et al*., 2014. The lysis buffer consisted of 10mM tris, 1mM EDTA, 1.0 % SDS, pH 8.0, and 20 mg/mL lysozyme (Sigma Aldrich, St. Louis, MO) (Adebayo et al., 2020; Adebayo et al., 2017; Kramer et al., 2018; Pearce et al., 2014). The urine pellet and lysis buffer were then incubated in a 37 °C water bath for 1 hour. The remainder of the extraction protocol was followed per manufacturer instructions with one modification. In the final step, DNA was eluted in 50 μl of elution buffer (AE), incubated at room temperature for five minutes; then, the eluent was run through the silica membrane of the spin column a second time.

### Magnetic Beads

Per Liu *et al*. (2017), urine pellets were resuspended in a lysis buffer composed of 10mM tris, 2mM EDTA, and 1% SDS, pH 8.0. The suspension was then frozen in liquid nitrogen for 1 minute followed by incubation in a 65 °C water bath for 5 minutes; the freeze / thaw process was repeated three times. After the third freeze / thaw step, suspensions were incubated for 30 minutes in a 65 °C water bath. Suspensions were then centrifuged at 20,000 x *g* for five minutes. After completing the lysis step and centrifugation, the supernatant was placed in PCR tube strips containing AMPure XP magnetic beads (Beckman Coulter, Indianapolis, IN). The supernatant and magnetic beads were homogenized and incubated at room temperature, then placed on a magnetic separator for 5 minutes. During this step, lysed DNA was drawn to magnetic beads in the tube strips. The remaining supernatant was removed and beads were washed with 80% ethanol. This was repeated twice. After washing, DNA-bound beads were dried in a 37 °C heat block for 15 minutes. Dried DNA-bound beads were then resuspended in 40 μl Qiagen^©^ C6 solution. Resuspended samples were then placed on a magnetic separator for 1 – 2 minutes to pellet beads. The resulting supernatant contained DNA used in downstream analyses.

### PowerFecal

Urine pellets were resuspended in lysis buffer (PowerBead Solution + C1) and subjected to two rounds of bead beating (6 m/s for 60 seconds with a five-minute rest between cycles). Samples then underwent multiple inhibitor removal and purification steps. The remainder of the extraction protocol was followed with two modifications. During the second round of centrifuging, after applying the ethanol-based wash solution (C5) to the spin column, samples were centrifuged for 2 minutes (instead of 1 minute) to remove residual wash solution. In the final step, DNA was eluted in 50 μl of elution buffer (C6), incubated at room temperature for five minutes; then, the eluent was run through the silica membrane of the spin column a second time.

### PowerFecal Pro

Urine pellets were resuspended in lysis buffer (CD1) and subjected to two rounds of bead beating (6 m/s for 60 seconds with a five-minute rest between cycles). Samples then underwent multiple inhibitor removal and purification steps. The remainder of the extraction protocol was followed with one modification. In the final step, DNA was eluted in 50 μl of elution buffer (C6), incubated at room temperature for five minutes; then, the eluent was run through the silica membrane of the spin column a second time.

DNA yields from all samples were then quantified on a Qubit^®^ 4.0 Fluorometer (Invitrogen, Thermo Fisher Scientific™, Carlsbad, CA, USA) using a 1X dsDNA High Sensitivity Assay. Hereafter, DNA concentrations measured using Qubit^®^ are referred to as total DNA concentrations.

### Quantification of bacterial DNA by qPCR

Bacterial DNA was amplified using 16S rRNA bacterial primers and probes per Nadkarni *et al*. (2002) on a QuantStudio™ 3 Real-Time PCR System (Applied Biosystems™, Thermo Fisher Scientific™, Carlsbad, CA, USA). 300 nM of forward primer (5’ – TCCTACGGGAGGCAGCAGT – 3’), 300 nM of reverse primer (5’ – GGACTACCAGGGTATCTAATCCTGTT – 3’), and 175 nM of probe ((6FAM) – 5’ – CGTATTACCGCGGCTGCTGGCAC – 3’ – (TAMRA)) were added to each reaction. qPCR cycling parameters were as follows: 50 °C for 2 min, 95 °C for 10 min (initial denaturation) and 40 cycles of 95 °C for 15 s (denaturation) and 60 °C for 1 min (annealing and extension) (Nadkarni et al., 2002). To be included in analysis, at least two replicates per sample had to amplify. Following qPCR, cycle thresholds were log_10_-transformed using the equation listed below under “qPCR standard curve,” and the antilog of each sample was used to calculate the bacterial DNA concentration in each sample.

### qPCR standard curve

DNA extracted from an *Escherichia coli* isolate was used to generate a standard curve for qPCR. Five ten-fold dilutions of *E. coli* DNA ranging from approximately 5 × 10^4^ pg/μl to 5 × 10^−1^ pg/μl were run in triplicate using the primers, probe, cycling parameters, and the QuantStudio instrument described above. DNA concentrations were then log_10_-transformed and plotted against cycle threshold values on a linear regression using R package ggplot2 v.3.3.2. The resulting equation was y = −5.329x + 36.504 (R^2^ = 0.984) where y is the cycle threshold and x is the log_10_-transformed DNA concentration. To calculate absolute cell counts in each sample, *Escherichia coli* was used as a standard, and the bacterial DNA concentrations were divided by the theoretical weight of one *E. coli* cell (4.96 fg DNA) per Nadkarni et al. 2002. Absolute cell counts were then multiplied by relative abundances of specific taxa (e.g. *Sphingomonas* and *Pasteurellaceae* bacterium canine oral taxon 272) to obtain and compare absolute cell counts of these taxa.

### 16S rRNA sequencing and sequence processing

DNA underwent library preparation and sequencing at Argonne National Laboratory. Library preparation was performed as follows: the V4 region of the 16S rRNA gene was amplified using primers 515F and 806R (Caporaso et al., 2012, 2011). PCR reactions (25 µL) contained 9.5 µL of MO BIO PCR Water (Certified DNA-Free), 12.5 µL of QuantaBio’s AccuStart II PCR ToughMix (2x concentration, 1x final), 1 µL Golay barcode tagged Forward Primer (5 µM concentration, 200 pM final), 1 µL Reverse Primer (5 µM concentration, 200 pM final), and 1 µL of template DNA. The following PCR conditions were applied: 94 °C for 3 minutes to denature the DNA, with 35 cycles at 94 °C for 45 s, 50 °C for 60 s, and 72 °C for 90 s; with a final extension of 10 min at 72 °C. Amplicons were then quantified using PicoGreen (Invitrogen) and a plate reader (Infinite→ 200 PRO, Tecan). Equimolar volumes of the amplicons were pooled into a single tube. This pool was then cleaned using AMPure XP Beads (Beckman Coulter), and quantified with a fluorometer (Qubit, Invitrogen). After quantification, the molarity of the pool was diluted to 2 nM, denatured, and then diluted to a final concentration of 6.75 pM with a 10% PhiX spike. Amplicons were sequenced on a 251bp x 12bp x 251bp Illumina MiSeq (Lemont, IL, USA) run using customized sequencing primers and procedures (Caporaso et al., 2012). Sequencing data is available at NCBI Bioproject PRJNA689589.

Raw, paired-end sequence reads were processed using QIIME2 v. 2020.2 (Boylen et al., 2019). The DADA2 plugin was used to truncate reads at 230 bp and trim 33 bp from the left side of both forward and reverse reads (Callahan et al., 2016). These parameters were used to ensure primers and barcodes were removed and to denoise paired end reads. Taxonomy was assigned in QIIME2 using the Silva 132 99% Operational Taxonomic Units (OTUs) from the 515F / 806R classifier (Quast et al., 2013; Yilmaz et al., 2014). We opted not to rarefy at 300 reads as this drastically decreased diversity and eliminated rare OTUs. Rarefaction can also increase Type 1 errors and variance where overdispersion can mask differential abundance between samples (McMurdie and Holmes, 2014). To avoid these issues and account for rare OTUs, an unrarefied table was used as input in α- and β-diversity metrics.

### Statistical analyses

Total DNA concentrations (as measured by Qubit) and bacterial DNA concentrations (as calculated from qPCR) were tested for normality using the Shapiro Wilk Normality Test in R version 3.5.2. DNA concentrations and 16S rRNA read numbers were analyzed using Kruskal-Wallis Rank Sum Test. Statistical significance was achieved if the p-value was less than 0.05.

All diversity metrics were computed using the R package phyloseq with a p-value cutoff of 0.05 adjusted using the Benjamini & Hochberg False Discovery Rates (McMurdie and Holmes, 2013). To analyze microbial diversity, three α-diversity metrics were used: Observed Operational Taxonomic Units (OTUs) (equivalent to richness), Shannon, and Simpson. Kruskal-Wallis Rank Sum Tests were used to compare α-diversity results to categorical variables of interest (extraction method, sex, dog). Post-hoc pairwise comparisons were calculated using Pairwise Wilcoxon Rank Sum Tests. To compare microbial composition between groups (extraction method, sex, dog), three β-diversity metrics were used: Bray Curtis, Unweighted UniFrac, and Weighted UniFrac. A permutational analysis of variance (PERMANOVA) based on Euclidean distance matrices and 1000 permutations was used to quantify these differences (adonis2 function, R package vegan v.2.5.6). Multilevel pairwise comparisons for dog and extraction method were calculated using pairwise PERMANOVAs with 1000 permutations (Martinez Arbizu, 2020). An Analysis of Composition in Microbiomes (ANCOM) in QIIME2 was used to identify differentially abundant taxa between groups (extraction method, sex, dog).

## Results

### Urine total DNA concentrations

Total DNA concentrations were measured on a Qubit fluorometer (**Table S2**) and were not normally distributed (Shapiro-Wilk Normality Test, *p* = < 0.0001). Twenty-six samples (9 PFP, 6 MB, 4 B, 4 PF, 3 BTL) and all 5 negative controls (one per method) had concentrations that were too low to read (< 0.01 ng/μl). Quantifiable DNA concentrations ranged from 0.02 ng/μl to 1.37 ng/μl for all other samples including positive controls. The number of samples with quantifiable DNA varied by extraction method (**Fig. 1a**). BTL extracted quantifiable DNA from eight out of 12 urine samples (including the positive control). B and PF extracted DNA from seven out of 12 urine samples (including the positive control). MB and PFP extracted DNA from five and two samples, respectively (**Fig. 1a**). Notably, PFP failed to extract DNA from the spiked positive control sample. Total DNA concentrations did not differ significantly by method (**Fig. 1b**, Kruskal-Wallis, *p* = 0.165), but did differ significantly by sex (**Fig. 1c**, Kruskal-Wallis, *p* = 0.0007) and by dog (**Fig. 1d**, Kruskal-Wallis, *p* = 0.001). Males had significantly higher DNA concentrations; however, no pairwise comparisons of DNA concentrations were significant by dog (**Table S3**). Bacteremia (B) produced the highest average total DNA concentrations, followed by PF, BTL, MB, and PF.

**Figure 1:**
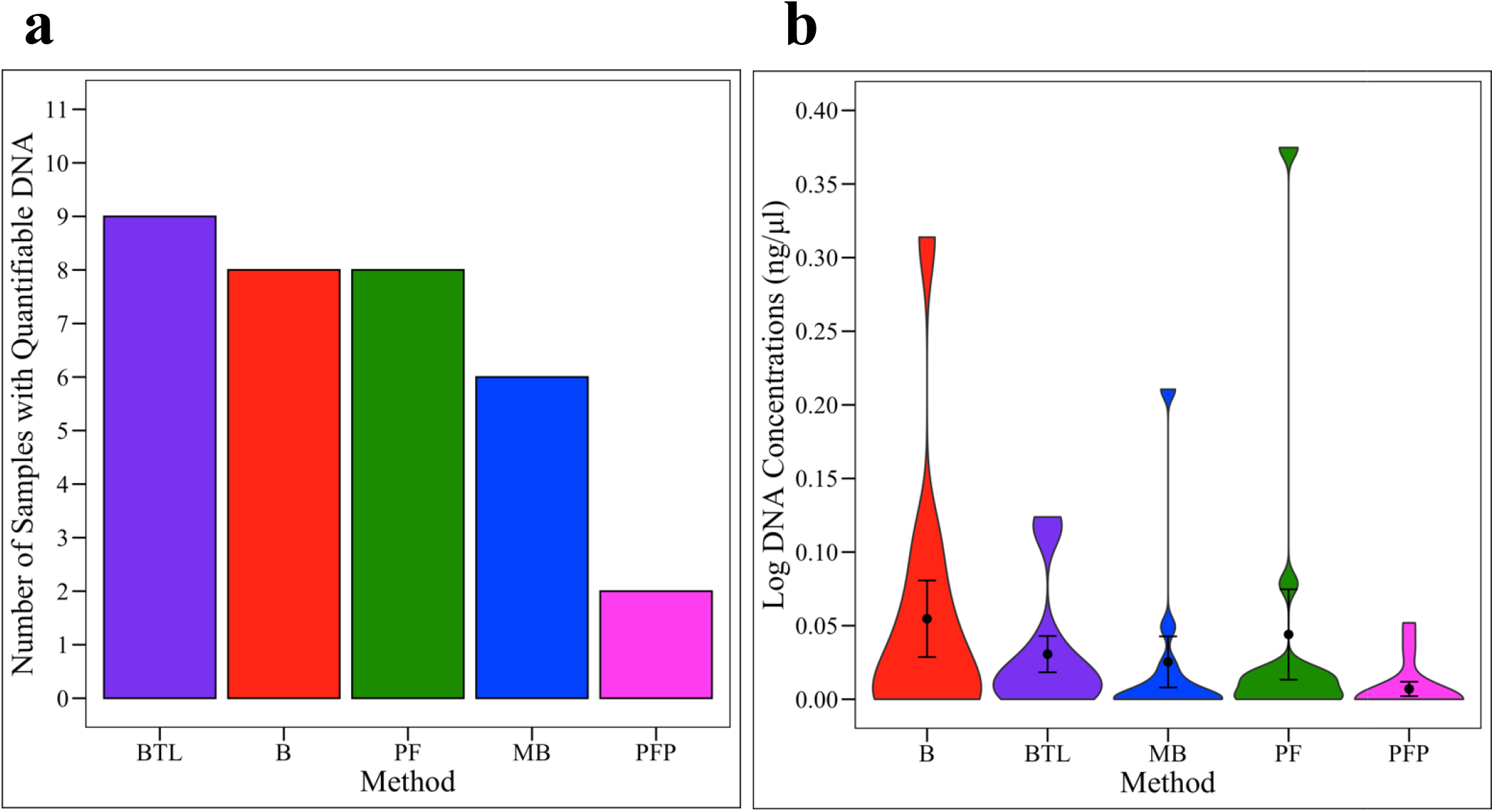
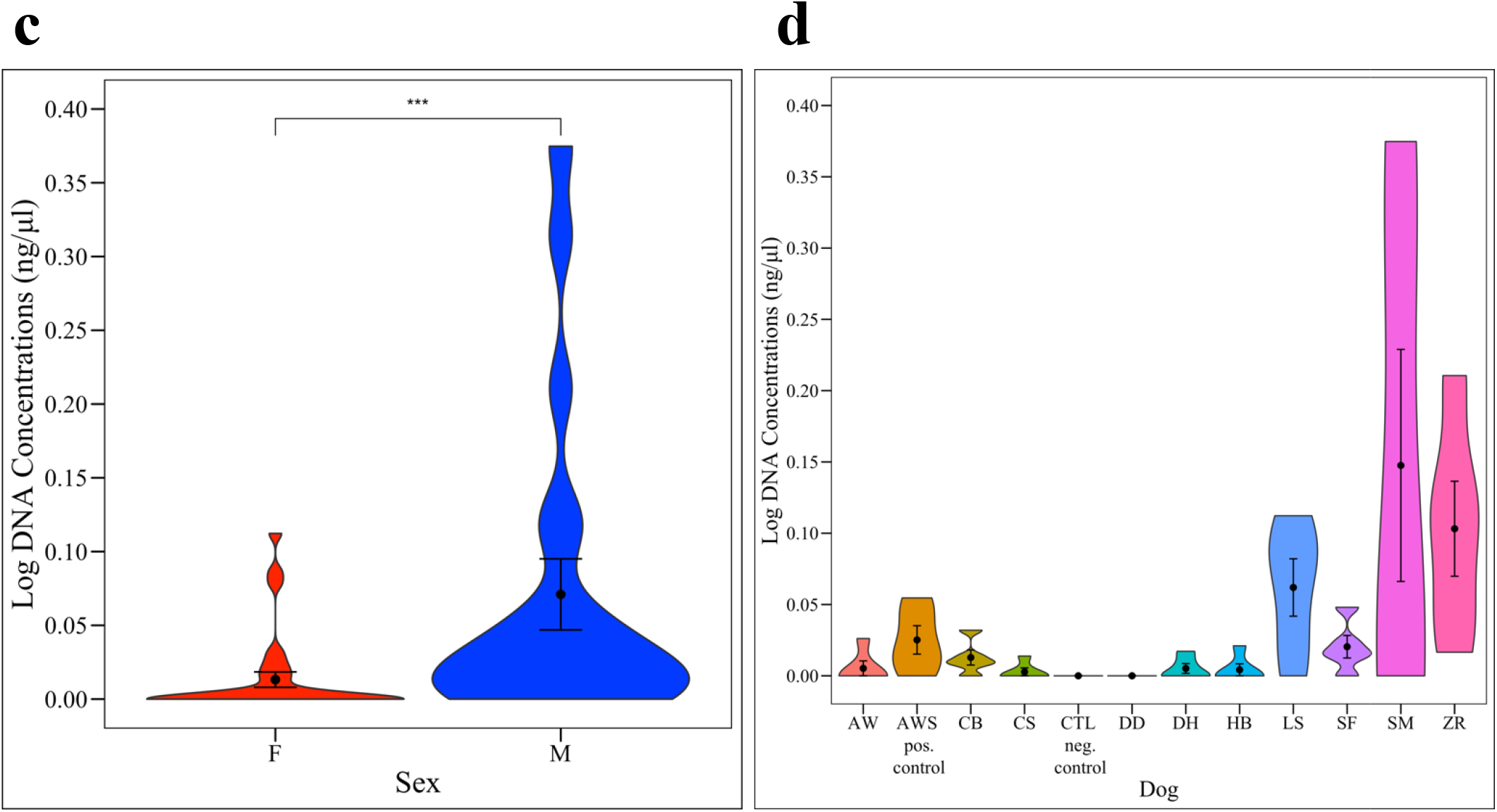
Total DNA concentrations. (**a**) Number of samples with measurable total DNA by extraction method (negative control excluded). Total DNA concentration, measured via Qubit, by (**b**) extraction method, (**c**) sex, and (**d**) dog. Total DNA concentrations did not differ significantly by extraction method (Kruskal-Wallis, *p* = 0.165) but did differ significantly by sex (Kruskal-Wallis, *p* = 0.0007; males > females), and dog (Kruskal-Wallis, *p* = 0.001). By dog, no pairwise comparisons were statistically significant. B = Bacteremia, BTL = Blood & Tissue with Lysozyme, MB = Magnetic Beads, PF = PowerFecal, PFP = PowerFecal Pro, F = Female, M = Male.

### Urine bacterial DNA concentrations

Bacterial DNA concentrations were measured in triplicate via qPCR (**Table S2)** and were not normally distributed (Shapiro-Wilk Normality Test, *p* = < 0.0001). Twenty-two samples (5 MB, 8 BTL, 5 PFP, 3 PF, 1 B) and all 5 negative controls did not amplify any bacterial DNA. Five samples (AWMB, LSMB, SMMB, SFMB, SMPFP) were excluded for failing to have at least two replicates amplify. All samples exhibited less than 3% variation in cycle threshold values between replicates with three exceptions: AWSMB (the spiked positive control, 5.6% variation), SFB (6.1%), and SMB (3.4%). All three samples were included in analyses. Quantifiable bacterial DNA concentrations ranged from 0.28 pg/μl to 729.48 pg/μl. The number of samples with quantifiable bacterial DNA varied by extraction method (**Fig. 2a**). Additionally, bacterial DNA concentrations differed significantly by method (**Fig. 2b**, Kruskal-Wallis, *p* = 0.044) and by dog (**Fig. 2d**, Kruskal-Wallis, *p* = 0.0005); although, no pairwise comparisons by method or dog were significant (**Table S4**). Bacteremia (B) yielded the highest bacterial DNA concentrations and extracted quantifiable bacterial DNA from the greatest number (10 out of 11) of urine samples. Bacterial concentrations did not differ significantly by sex (**Fig. 2c**, Kruskal-Wallis Rank Sum Test, *p* = 0.333). Bacterial and total DNA concentrations were significantly correlated (**Fig. S1a**, R = 0.42, *p* = < 0.001), and 13 samples were identified as having greater bacterial DNA concentrations than total DNA concentrations (**Table S2**).

**Figure 2:**
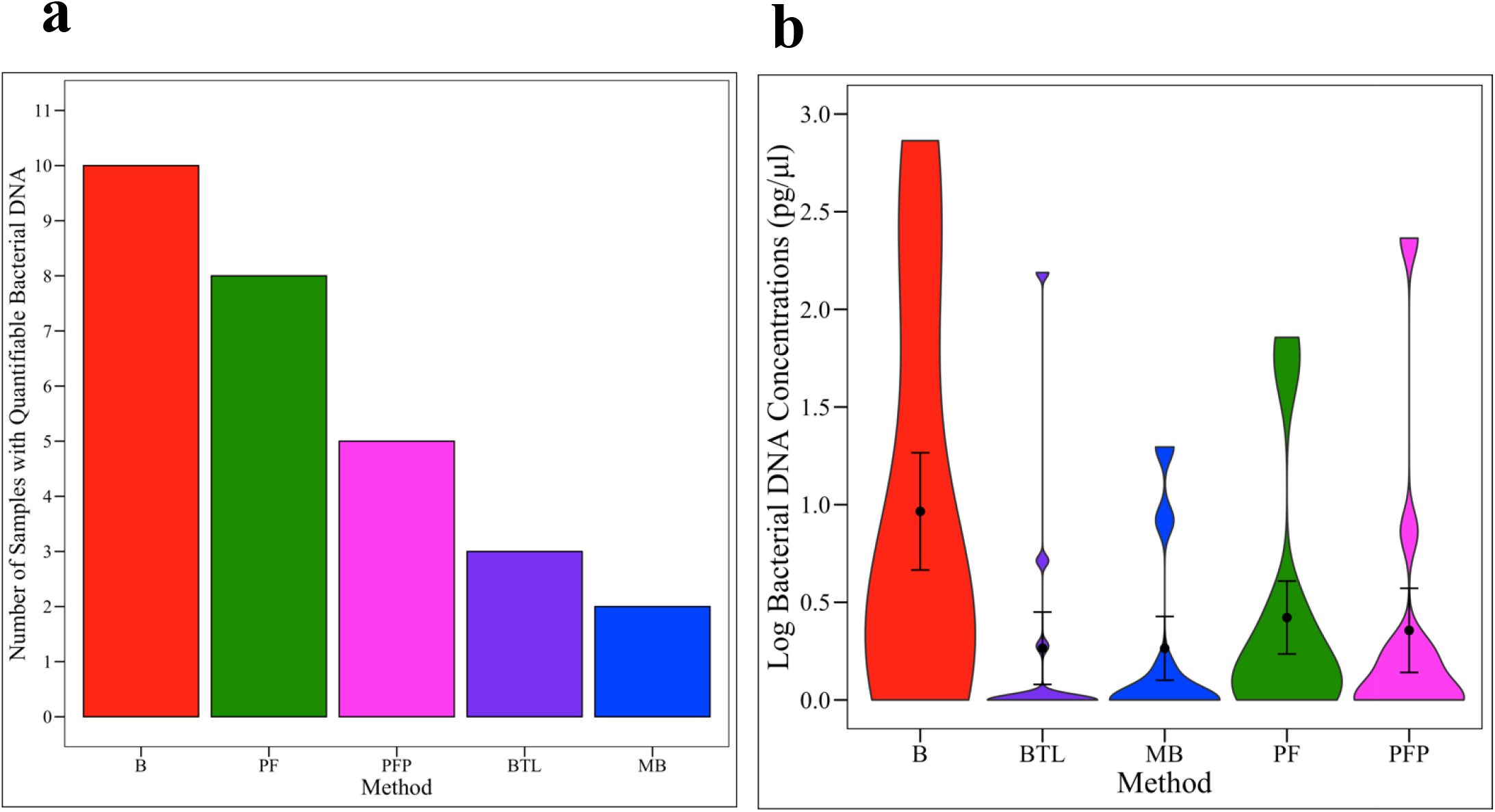
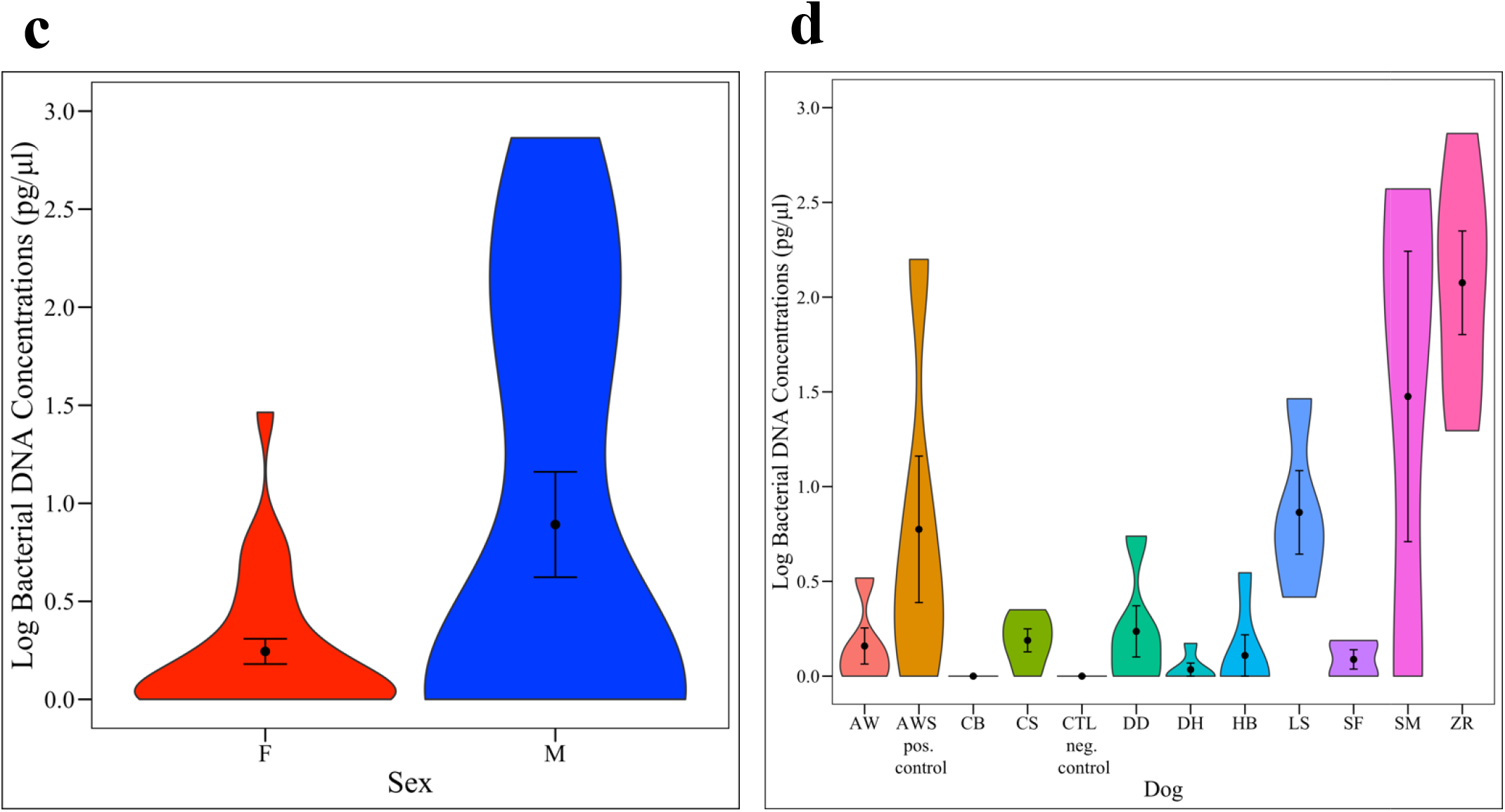
Bacterial DNA Concentrations. (**a**) Number of samples with measurable bacterial DNA by extraction method. Bacterial DNA concentrations, calculated via qPCR, by (**b**) extraction method, (**c**) sex and (**d**) dog. Bacterial DNA concentrations differed significantly by extraction method (Kruskal-Wallis, *p* = 0.044) and by dog (Kruskal-Wallis, *p* = 0.0005); although, no pairwise comparisons were significant. Bacterial DNA concentrations did not differ significantly by sex (Kruskal-Wallis, *p* = 0.333; positive control excluded). B = Bacteremia, BTL = Blood & Tissue with Lysozyme, MB = Magnetic Beads, PF = PowerFecal, PFP = PowerFecal Pro, F = Female, M = Male.

### Microbial diversity by extraction method, dog, and sex

Based on the composition of the negative controls, the following taxa were deemed to be contaminants and were bioinformatically removed from all samples: *Bradyrhizobium*, Caulobacteraceae, Chloroflexi, Cyanobacteria, *Micrococcus*, and *Prevotella 9*. Chloroplasts, mitochondria, and any reads identified as Eukarya or Archaea were also removed from all samples. Additionally, four samples (CBMB, SFBTL, CBBTL, CBPFP) with fewer than 300 reads were excluded from 16S rRNA bacterial community analyses. Three of these four samples came from dog CB, who had one of the lowest urine bacterial concentrations. Three of the 5 negative controls also had fewer than 300 reads while the remaining 2 negative controls had 8101 and 8269 reads respectively and a taxonomic composition suggesting potential cross-contamination by urine microbiota during the plating / library preparation / sequencing process. The remaining samples ranged from 316 to 42,090 reads (average = 16,126). A total of 51 samples (11 B and PF, 10 MB and PFP, 9 BTL) were retained for analysis. The number of 16S rRNA reads per sample differed significantly by dog (Kruskal-Wallis, *p* = <0.00001), but not by sex (Kruskal-Wallis, *p* = 0.937) or extraction method (Kruskal-Wallis, *p* = 0.378); although, Bacteremia yielded the greatest number of 16S reads per sample (**Fig. S2**,). There was also a significant correlation between the number of 16S reads and bacterial DNA concentrations (**Fig. S1b**, R = 0.28, *p* = 0.047).

Microbial diversity was compared across samples by extraction method, dog, and sex (Shannon Index: **Fig. 3**; Observed OTUs and Simpson Index: **Fig. S3)**. The positive control, which was urine from female dog AW spiked with *M. plutonius*, was removed from all analyses by sex to prevent bias. All three measures of diversity revealed the same patterns. Microbial diversity did not differ by extraction method (Kruskal-Wallis: Shannon, *p* = 0.95; Observed OTUs, *p* = 0.727, Simpson, *p* = 0.958; **Fig. 3a, S3a**,**d**) but did differ significantly by sex (Kruskal-Wallis: Shannon, *p* = 0.00005; Observed OTUs, *p* = 0.002, Simpson, *p* = 0.0001; **Fig. 3b, S3b**,**e)** and by dog (Kruskal-Wallis: Shannon, *p* = 0.00002; Observed OTUs, *p* = 0.0001, Simpson, *p* = 0.00002; **Fig. 3c, S3c**,**f, Table S5)**. Females had significantly higher microbial diversity than males across all diversity metrics. By dog, LS had significantly higher microbial diversity than 9 other dogs while ZR had significantly lower microbial diversity than 9 other dogs. For a full list of significant pairwise comparisons by dog, see **Table S5**.

**Figure 3.**
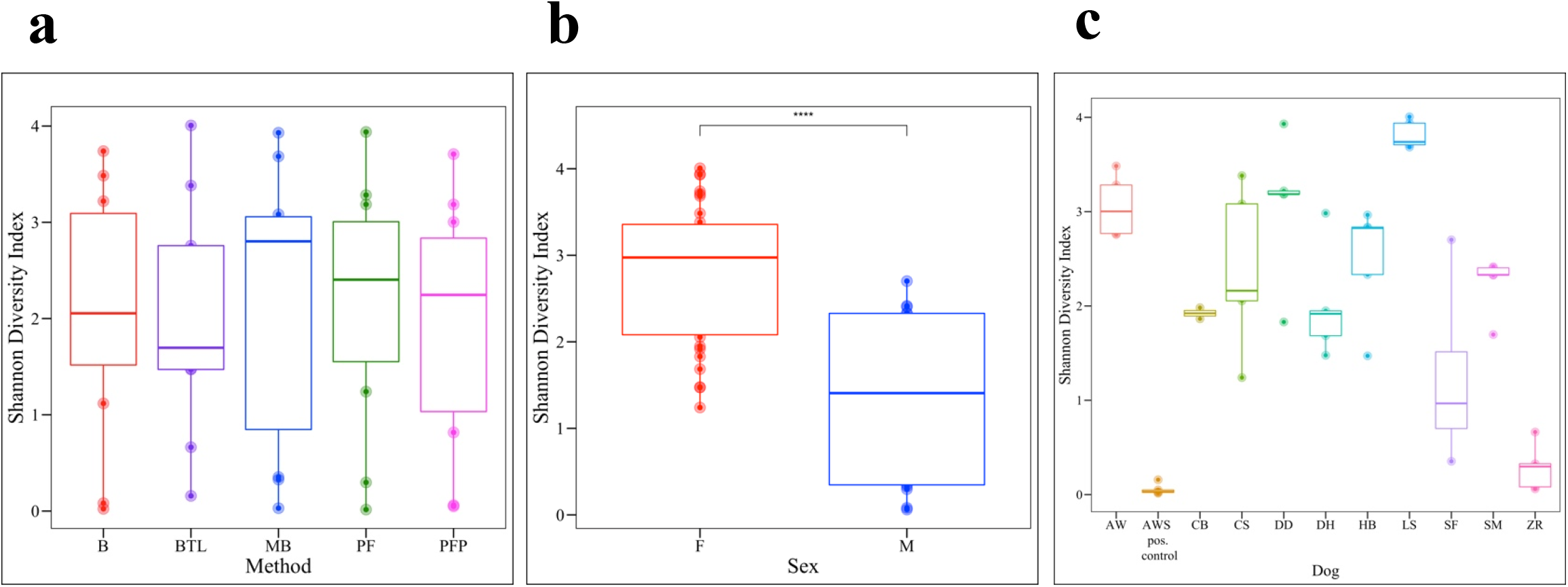
Microbial diversity. The Shannon diversity metric was used to compare microbial diversity by (**a**) extraction method, (**b**) sex, (**c**) and dog. Microbial diversity did not differ significantly by kit (Kruskal-Wallis, *p* = 0.95) but did differ significantly by dog (Kruskal-Wallis, *p* = 0.00002). For all statistically significant pairwise comparisons by dog, see **Table S3**. Females exhibited higher microbial diversity than males (Kruskal-Wallis, *p* = 0.00005). B = Bacteremia, BTL = Blood & Tissue with Lysozyme, MB = Magnetic Beads, PF = PowerFecal, PFP = PowerFecal Pro, F = Female, M = Male

### Microbial composition by extraction method, dog, and sex

Bray Curtis (**Fig. 4**) and Unweighted and Weighted UniFrac metrics (**Fig. S4**) were used to compare microbial composition (beta-diversity) across groups. No significant differences were observed by extraction method (PERMANOVA: Bray Curtis, *p* = 0.999; Unweighted UniFrac, *p* = 0.478; Weighted UniFrac, *p* = 0.524, **Fig. 4a, Fig. S4a**,**d**). However, microbial composition did differ significantly by sex (PERMANOVA: Bray Curtis, *p* = < 0.001, Unweighted UniFrac, *p* = 0.005; Weighted UniFrac, *p* = 0.011 **Fig. 4b, S4b**,**e**) and by dog (PERMANOVA: Bray Curtis, *p* = < 0.001, Unweighted UniFrac, *p* = 0.005; Weighted UniFrac, *p* = 0.011; **Fig. 4c, S4c**,**f, Table S6**).

**Figure 4.**
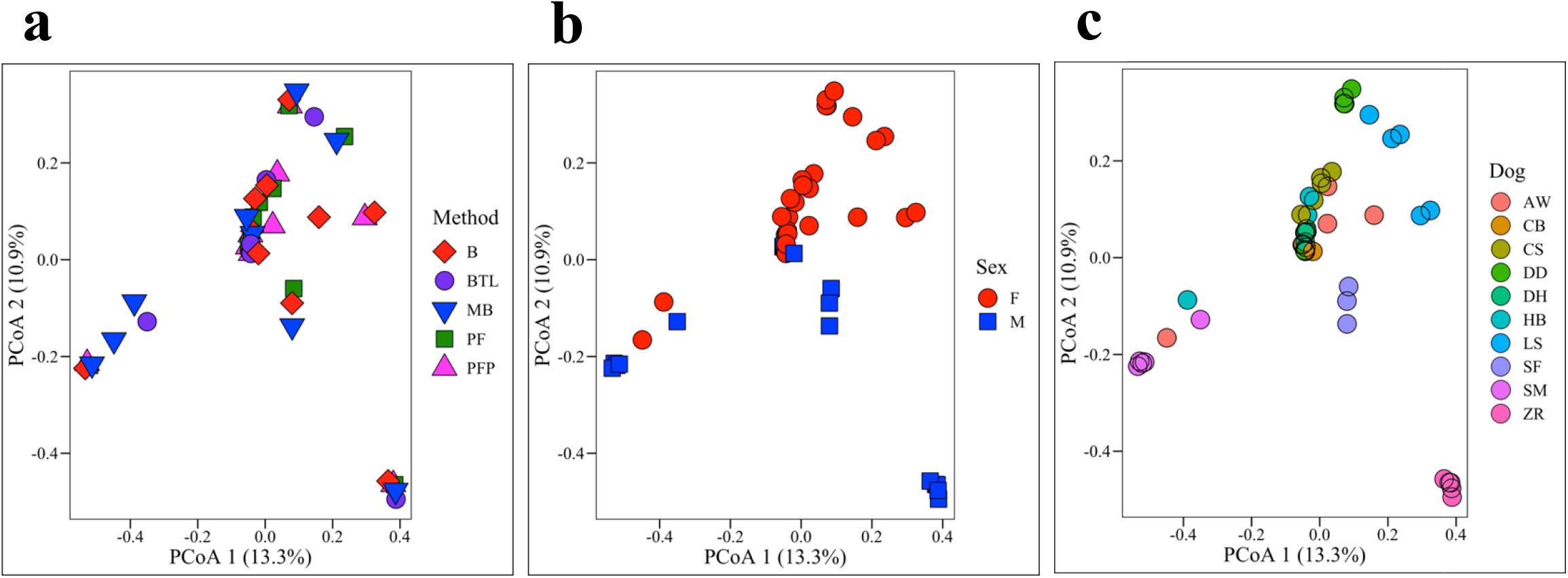
Microbial Composition. Bray Curtis dissimilarity matrices were used to compare microbial composition (beta-diversity) by (**a**) extraction method, (**b**) sex, (**c**) and dog. Microbial composition did not differ significantly by method (PERMANOVA, *p* = 0.999) but did differ significantly by sex (PERMANOVA, *p* = < 0.001) and dog (PERMANOVA, *p* = < 0.001).

### Bacterial taxonomic differences by extraction method, dog, and sex

In total, there were 21 phyla, 323 genera, and 203 amplicon sequence variants (ASVs, roughly equivalent to species) observed across all canine urine samples. Collectively, the three most abundant phyla across all samples were Proteobacteria, Bacteroidetes, and Firmicutes (**Fig. 5**). We also noted that the positive control sample, AWS, which was urine from dog AW spiked with *M. plutonius*, had significantly lower microbial diversity (Shannon, Wilcoxon Rank Sum test, *p* = 0.022) as compared to AW. This indicates that in urine dominated by a specific microbe (e.g. during a urinary tract infection), amplicon sequencing precludes the ability to detect other microbes present in the microbial community. To test for differentially abundant taxa between groups, we first removed all taxa present at < 1% relative abundance. At the phyla level, there were no significant differences in taxa abundances by extraction method (**Fig. 5a**, Kruskal-Wallis, *p* = 0.81) or dog (**Fig. 5c**, Kruskal-Wallis, *p* = 0.12), but females had a significantly higher abundance of Actinobacteria as compared to males (**Fig. 5b**, ANCOM, *W* = 14). At the L7 (roughly species) level, no taxa were found to be differentially abundant by extraction method. However, several taxa were identified as differentially abundant by sex and by dog. By sex, the relative abundance of *Sphingomonas* was significantly increased in females compared to males (**Fig. S5a**, ANCOM *W* = 573) while *Pasteurellaceae* bacterium canine oral taxon 272 was significantly more abundant in males (**Fig. S5b**, ANCOM *W* = 591). Thirty-seven taxa (L7 level) were differentially abundant by dog (**Table S7**, ANCOM).

**Figure 5.**
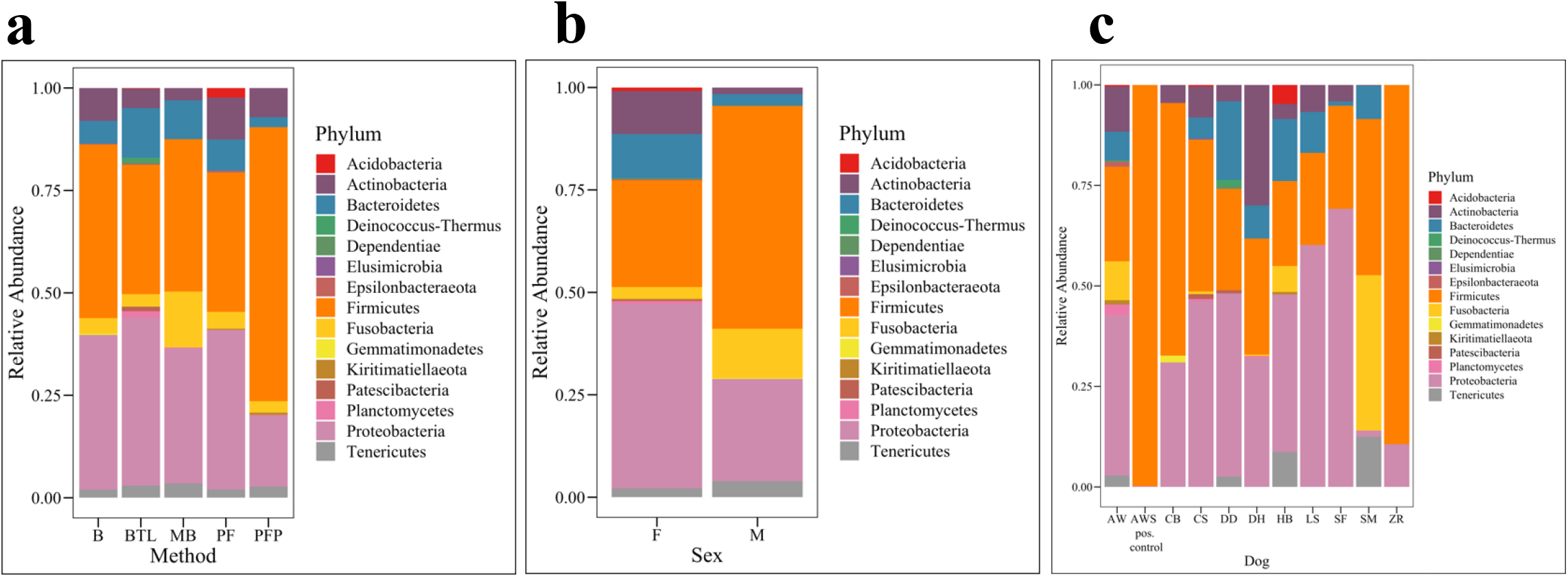
Bacterial Relative Abundances by Phyla. Microbial taxa bar plots by Phyla for (**a)** extraction method, (**b)** sex and (**c)** dog. B = Bacteremia, BTL = Blood Tissue with Lysozyme, MB = Magnetic Beads, PF = PowerFecal, PFP = PowerFecal Pro, F = Female, M = Male.

We then converted the relative abundances into absolute cell counts using bacterial DNA concentrations derived from qPCR and using the weight of DNA in one *E. coli* cell (4.96 fg) as a standard (Nadkarni et al., 2002). Total cell counts ranged from approximately 10 – 147,072 cells per sample. We multiplied total cell counts by the relative abundances of *Sphingomonas* or *Pasteurellaceae* bacterium canine oral taxon 272 to get the absolute cell counts of these taxa. The absolute cell counts of *Sphingomonas* were significantly increased in females (**Fig. 6a**, Kruskal-Wallis, *p* = 0.0148) while the absolute cell counts of *Pasteurellaceae* bacterium canine oral taxon 272 were significantly increased in males (**Fig. 6b**, Kruskal-Wallis, *p* = 0.0002).

**Figure 6.**
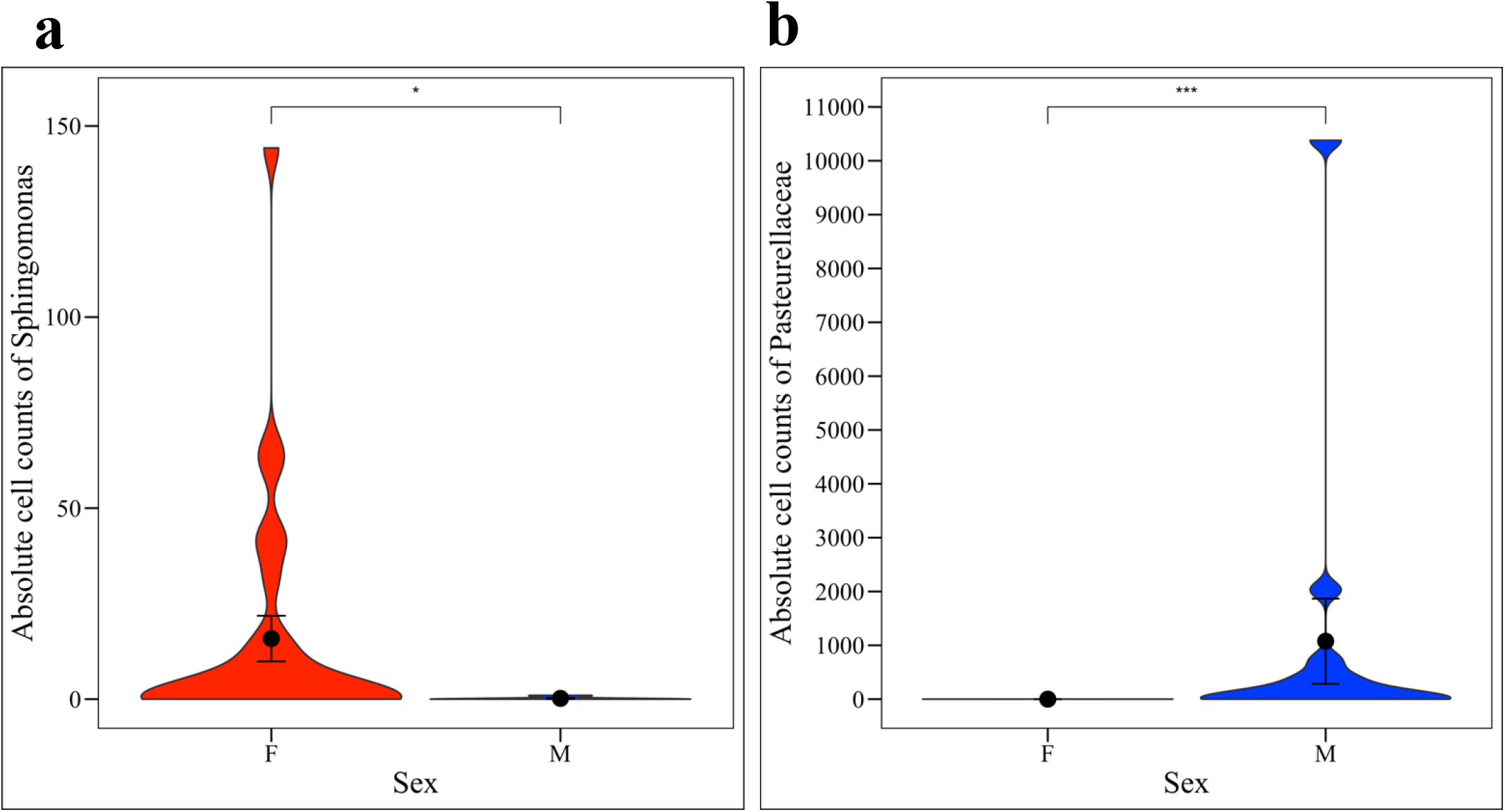
Differentially Abundant Taxa by Sex. (**a)** Females had significantly greater absolute cell counts of *Sphingomonas* (Kruskal-Wallis, p = 0.0148) while (**b)** males had significantly greater absolute cell counts of *Pasteurellaceae* bacterium canine oral taxon 272 (Kruskal-Wallis, p = 0.0002).

## Discussion

We compared total and bacterial DNA concentrations as well as 16S rRNA microbial community sequencing data from the urine of 10 healthy dogs extracted using 5 different DNA isolation methods. Each method employed various mechanical, chemical, and / or thermal lysing techniques. Sex and dog, but not extraction method, significantly affected DNA concentrations and microbial diversity and composition. Bacteremia (B) was determined to be one of the most effective methods for urine microbial DNA extraction.

### DNA concentrations and 16S reads

Bacteremia (B) extracted the greatest total (although not significant) and bacterial DNA concentrations from the canine urine samples (**Fig. 1b, 2b**). Moreover, Bacteremia extracted quantifiable *bacterial* DNA from the greatest number of samples (10 out of 11 samples – including the positive control) while BTL extracted quantifiable *total* DNA from the greatest number of samples (9 out of 11) (**Fig. 1a, 2a**). B and PF each extracted total DNA from the second greatest number of samples (8 out of 11). Males contained significantly greater total but not bacterial DNA as compared to females (**Fig. 1c, 2c**). This contrasts a previous study on human urine microbiota which reported higher total DNA concentrations in females as compared to males (El Bali et al., 2014). Total and bacterial DNA concentrations and number of 16S rRNA reads also varied significantly by dog (**Fig. 1d, 2d, S2c, Table S3, S4**). Bacteremia also produced the greatest number of 16S rRNA reads as compared to other extraction methods (Kruskal-Wallis, *p* = 0.378); however, as 16S rRNA sequencing data are compositional, we do not weigh this finding heavily (Gloor et al., 2017) (**Fig. S2a**). Taken together, Bacteremia demonstrated efficacy over other methods in extracting DNA from dog urine, while biological factors such as sex and dog had strong effects on DNA concentrations.

In 13 samples, bacterial DNA concentrations were greater than total DNA concentrations. This primarily occurred in 3 dogs (AW, DD, ZR) and could be due to relatively high bacterial loads in these dogs, as well as the increased sensitivity of qPCR (bacterial DNA concentrations) as compared to Qubit (total DNA concentrations) (Hussing et al., 2018). In 9 other samples, 5 of which were extracted by BTL, there was quantifiable total DNA but no quantifiable bacterial DNA present, suggesting that these samples may have contained more host than bacterial DNA, or that BTL was more effective in extracting host as compared to bacterial DNA. Despite the lack of quantifiable bacterial DNA in these samples, we obtained 16S rRNA sequencing reads from all 9 samples; although, 3 of the 10 samples were excluded from 16S rRNA analysis for having fewer than 300 reads. This indicates that 16S rRNA sequencing may be more sensitive to bacterial DNA than qPCR (Charlebois et al., 2020); however, the microbial taxa present in these samples should be reviewed carefully for potential contamination as they could contain low reads and skewed relative abundances. Alternately, the different primer sets used in qPCR and 16S rRNA sequencing, although both considered “universal bacterial primers,” may contribute to the differences we observed in bacterial DNA detection between qPCR and 16S rRNA sequencing.

### Microbial Diversity and Composition

Microbial diversity and composition differed significantly by sex and dog but not extraction method (**Fig. 3, 4**), indicating that individual differences in the urine microbiota overwhelmed potential differences introduced by extraction. Four samples were excluded from this analysis for having fewer than 300 reads. All 10 canine urine samples extracted via B and PF (excluding negative and positive controls) contained greater than 300 reads and were retained for analysis. MB and PFP extractions each retained 9 samples for analysis, and BTL retained 8 samples. These results again highlight Bacteremia as a viable extraction method for urine microbiota, as it did not obviously skew microbial communities while also generating reasonable 16S rRNA yields. We further observed that urine microbiota were highly variable between individuals, a finding that has been reported previously in studies on human and canine urine (Gottschick et al., 2017; Hilt et al., 2014; Pearce et al., 2014; Wolfe et al., 2012). A few dogs stood out in terms of microbial composition and diversity. ZR, for example, had significantly lower microbial diversity (Shannon) than 7 out of 9 dogs (**Table S5**) and significantly different microbial composition (Bray-Curtis) than 8 out of 9 dogs (**Table S6**). LS had significantly higher microbial diversity (Shannon) than 6 out of 9 dogs and significantly different microbial composition (Bray-Curtis) than 8 out of 9 dogs (**Table S5, S6**).

We also observed that females had significantly higher microbial diversity than males despite the finding that males had significantly higher total (**Fig. 1c**) and bacterial DNA (**Fig. 2c**); although, the latter was not significant. This suggests that males may be shedding more host cells into urine than females. The increased urine microbial diversity in females could be due to differing anatomy between sexes, differing hormone profiles, or differing urination habits. In humans, urine microbial diversity results vary by study. In a 2013 study based on free-catch urine, increased diversity was reported in healthy females as compared to males (Lewis et al., 2013), while in a study from 2020 that compared both free-catch and catheterized urine, no differences in microbial diversity were reported between males and females (Pohl et al., 2020). In the only study, to our knowledge, on healthy canine urine microbiota, microbial diversity did not differ by sex (Burton et al., 2017); although, this study used cystocentesis to collect urine, reducing the potential for genital and skin contaminants that may be present in free-catch samples, as collected in our study. Hormones have also been linked to changes in the fecal microbiome of women and could feasibly be altering the urine microbiota as well (Fuhrman et al., 2014). Urinary behavior also differs between male and female dogs with males generally urinating more frequently than females (Wirant and McGuire, 2004). It is feasible that urine volume and retention time in the bladder could alter urine composition and the urine microbial community.

### Taxonomic differences

In this study, the three most abundant phyla across all samples were Proteobacteria, Bacteroidetes, and Firmicutes. Other studies on urine microbiota in humans and in dogs report similar findings (Karstens et al., 2016; Lewis et al., 2013; Nelson et al., 2010; Pearce et al., 2014; Siddiqui et al., 2011). There were no differentially abundant taxa at the phyla level by extraction method or dog, but the relative abundance of Actinobacteria was significantly higher in females than males. Actinobacteria has also been reported in the urine of human females (Karstens et al., 2016; Lewis et al., 2013; Pearce et al., 2014; Siddiqui et al., 2011; Thomas-White et al., 2018), and in the oral (Oh et al., 2015) and gut (Honneffer et al., 2017; Kerr et al., 2013) microbiota of dogs. At the L7 (roughly species) level, we observed several differentially abundant taxa by sex and by dog, but not by extraction method. Notably, *Pasteurellaceae* bacterium canine oral taxon 272 was significantly increased in males while *Sphingomonas* was significantly increased in females. Taxa in the Pasteurellaceae family have been reported as part of the canine oral (Dewhirst et al., 2012; Oh et al., 2015; Ruparell et al., 2020), nasal (Tress et al., 2017), and gut microbiota (Xenoulis et al., 2008). It is possible that this taxon is introduced into canine urine through licking of the prepuce or penis. As such, this taxon could represent a skin contaminant or could be a true inhabitant of canine urine. Similarly, *Sphingomonas* has been reported in the canine vaginal microbiota (Burton et al., 2017) and could represent a genital contaminant or true inhabitant of urine. In humans, *Lactobacillus* species are common vaginal microbes, but studies on urine microbiota collected via catheter demonstrate that similar or identical *Lactobacillus* species are also present and culturable from the bladder and are not just contaminants (Jacobs et al., 2020; Komesu et al., 2020; Thomas-White et al., 2018).

There were several limitations to the work performed here. First, mid-stream free-catch urine was used for this study. This collection technique is highly relevant as it is non-invasive and commonly employed in canine health assessments; however, it is subject to contamination by urethral, genital, and skin microbiota. In a previous study on canine urine microbiota that collected urine via antepubic cystocentesis, no significant differences in microbial composition or diversity were observed between male and female dogs (Burton et al., 2017), while in our study, significant differences in microbial composition and diversity were observed by sex. These differences could be attributed to genital (e.g. vaginal) or skin contaminants in free-catch urine. A second limitation in this study: We cannot determine if the DNA and sequences detected in urine samples came from live or dead bacteria. In other words, we may be detecting microbes that are not actually contributing to the urogenital environment. Specialized culture or assessments of microbial function (e.g. metabolomics, proteomics, transcriptomics) are necessary to make this distinction. Finally, these samples were analyzed using 16S rRNA sequencing. We did this to ensure that we could obtain valid sequencing data from a relatively small amount of urine (3 ml) with generally low DNA concentrations. Now that we have established an effective method for urine microbiota extraction, we can pursue deeper sequencing (e.g. whole shotgun metagenomic sequencing) to more fully characterize microbial genomes and potential microbial function. Also worth noting, we used 3 ml of urine for DNA extractions. Other studies on the urine microbiota have used a range of urine volumes from 1 ml (Pohl et al., 2020; Price et al., 2020; Thomas-White et al., 2017) to 30 ml (Burton et al., 2017; Shrestha et al., 2018). We opted to test smaller volumes of urine as it is not always feasible to obtain 30 ml of urine from a dog, particularly a small dog. Larger volumes of urine may yield larger DNA concentrations and, as such, more readily facilitate deeper sequencing of these samples.

## Conclusion

The Bacteremia (B) kit yielded the highest total DNA concentrations, the highest bacterial DNA concentrations, the greatest number of 16S rRNA sequencing reads, and it extracted bacterial DNA from the greatest number of samples. Moreover, microbial diversity and composition did not significantly differ by kit indicating that no method, including Bacteremia, dramatically biased the sequencing results. As such, Bacteremia proved effective as an extraction method for studies of the urine microbiota.

## Author Contributions

RM contributed to the study design, performed qPCR, analyzed sequencing results, and drafted the manuscript.

CM performed DNA extractions, assisted with data analysis, and provided feedback on the manuscript.

ME assisted with processing of sequencing data and provided feedback on the manuscript. VLH designed and directed the study, and aided in analysis of sequencing results, data interpretation, and manuscript preparation.

## Acknowledgments

We are grateful to Dr. Sushmitha Durgam for the shared used of her qPCR instrument and to the many dogs and dog owners who participated in this study.

