## Supplemental Materials for "Evaluating Extraction Methods to Study Canine Urine Microbiota"

**Figure S1 – Correlation Analyses**

**a**

**b**

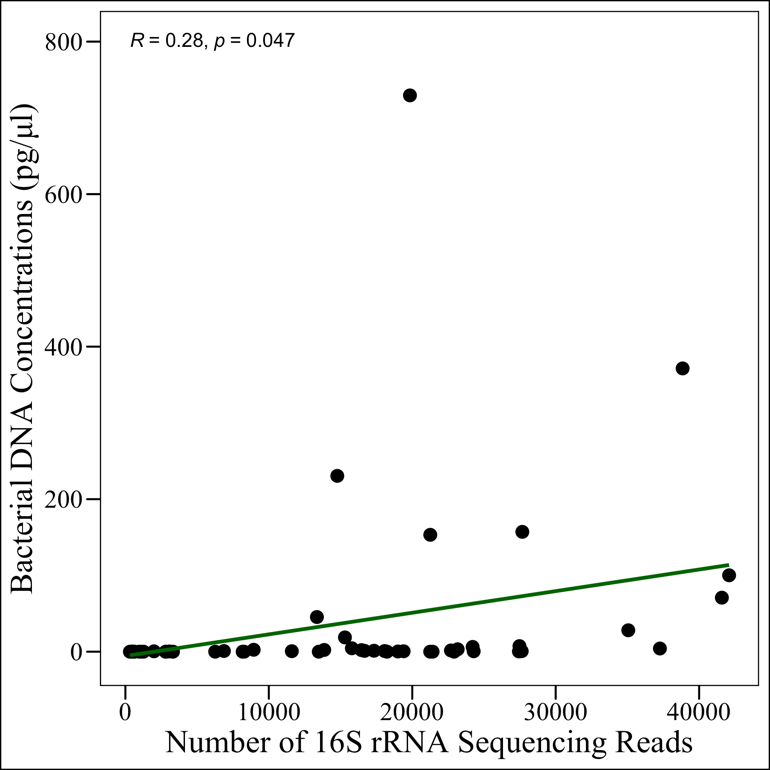

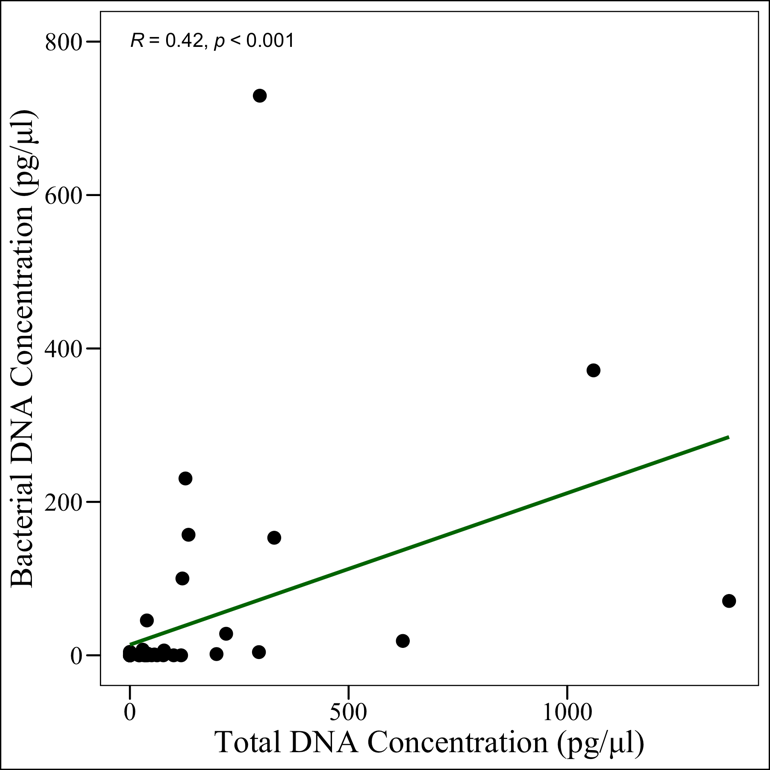

**Supplemental Figure 1 – Correlation Analyses.** Correlation between (**a)** total and bacterial DNA concentrations and (**b)** the number of 16S rRNA sequencing reads and bacterial DNA concentrations.

**Figure S2 – 16S rRNA Sequencing Reads**

**a**

**c**

**b**

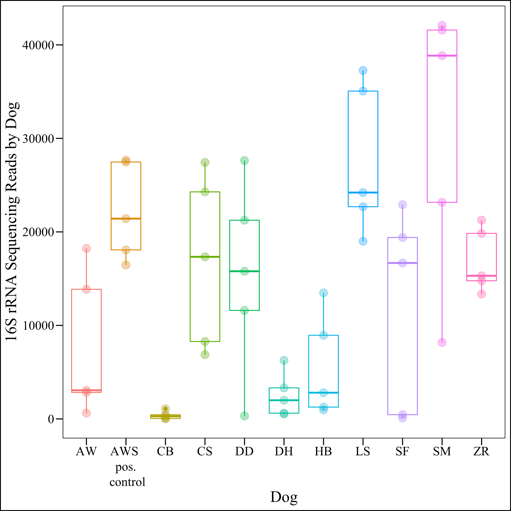

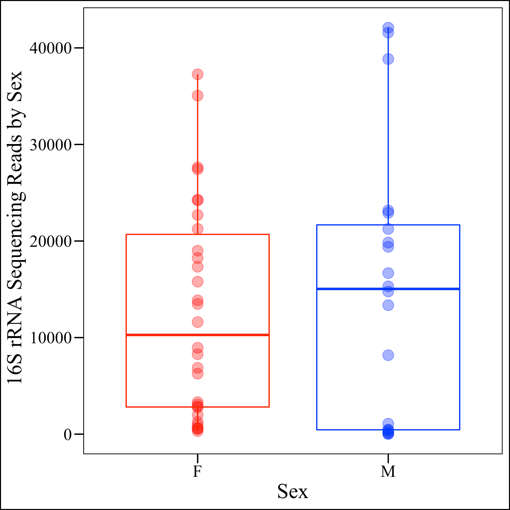

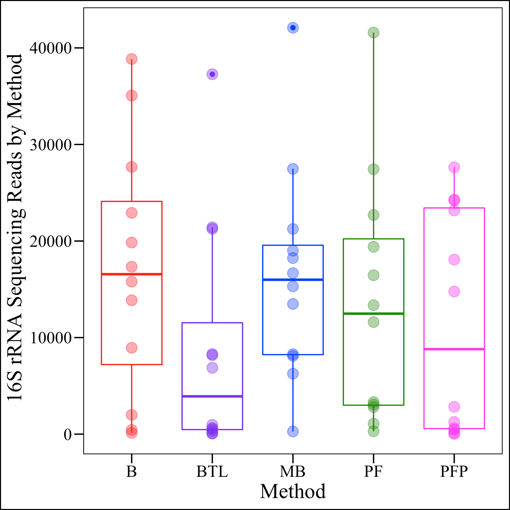

**Supplemental Figure 2 – 16S rRNA Sequencing Reads.** The number of 16S reads per sample was compared by (**a**) extraction method, (**b**) sex, and (**c**) dog. There were no significant differences in the number of reads by method (Kruskal-Wallis, *p* = 0.378) and sex (Kruskal-Wallis, *p* = 0.937) but there was a significant difference in the number of reads by dog (Kruskal-Wallis, *p* = <0.00001).

**Figure S3 – Microbial Diversity**

**a**

**b**

**c**

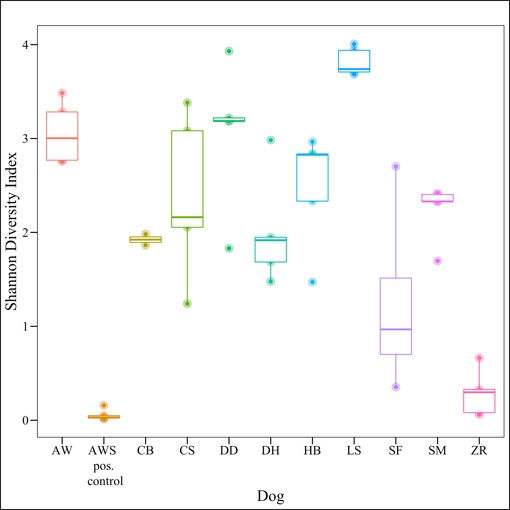

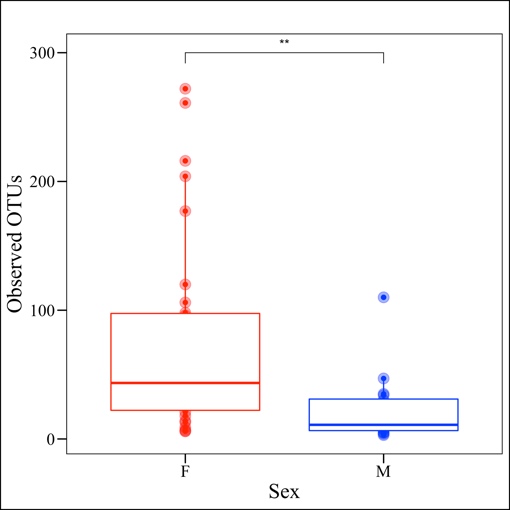

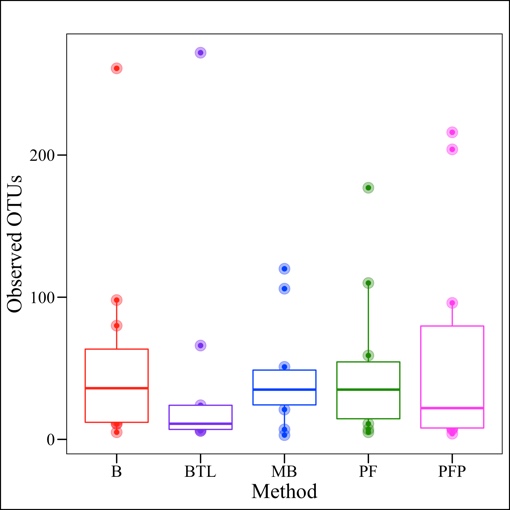

**f**

**e**

**d**

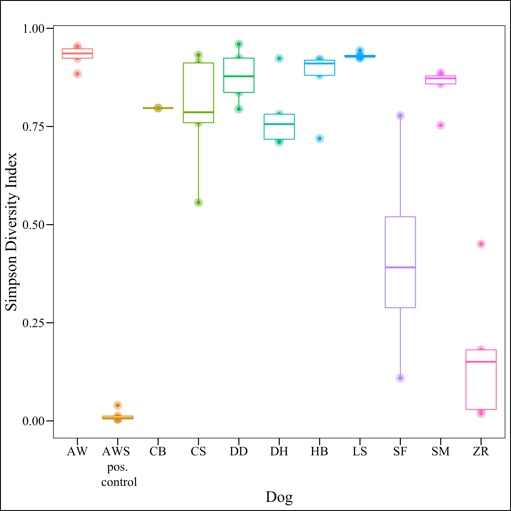

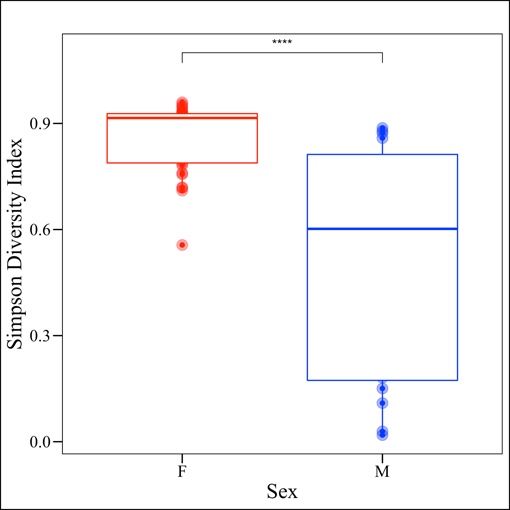

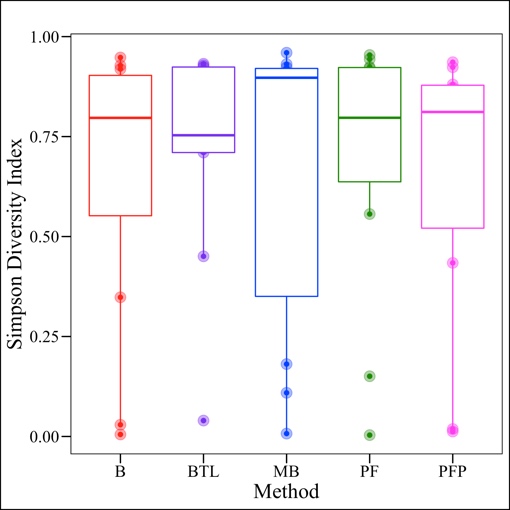

**Supplemental Figure 3 – Microbial Diversity.** Observed OTUs and the Simpson index were used to compare microbial diversity by extraction method (**a, d**), sex (**b, e**), and dog (**c, f).** Microbial diversity did not differ by extraction method (Kruskal-Wallis; Observed OTUs, *p* = 0.727; Simpson Index, *p* = 0.958) but did differ by dog (Kruskal-Wallis; Observed OTUs, *p* = 0.0003; Simpson Index, *p* = 0.00002). For all statistically significant pairwise comparisons by dog, see Supp. Table S3. Females exhibited significantly higher microbial diversity than males (Kruskal-Wallis: Observed OTUs, *p* = 0.002; Simpson, *p* = 0.0001). B = Bacteremia, BTL = Blood Tissue with Lysozyme, MB = Magnetic Beads, PF = PowerFecal, PFP = PowerFecal Pro, F = Female, M = Male.

**Figure S4 – Microbial Composition**

**c**

**b**

**a**

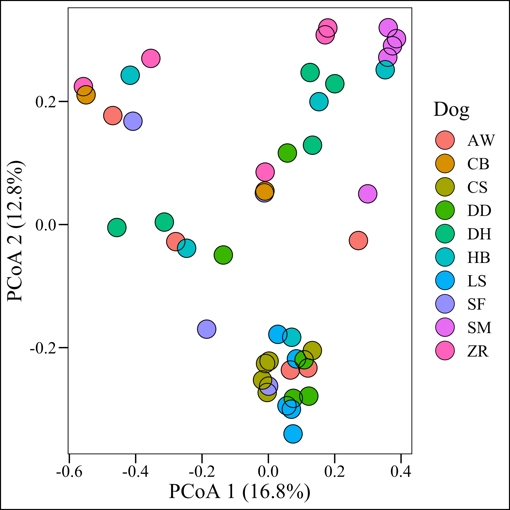

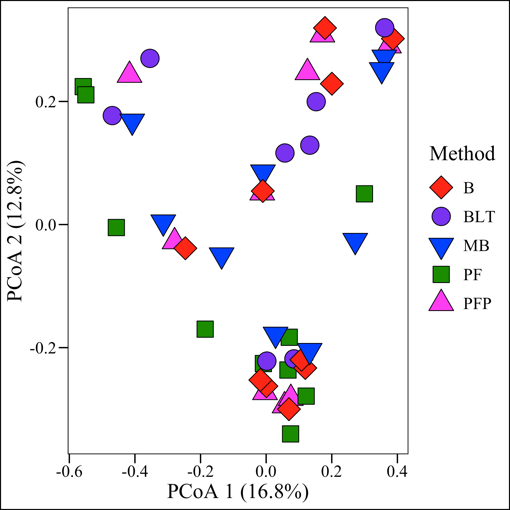

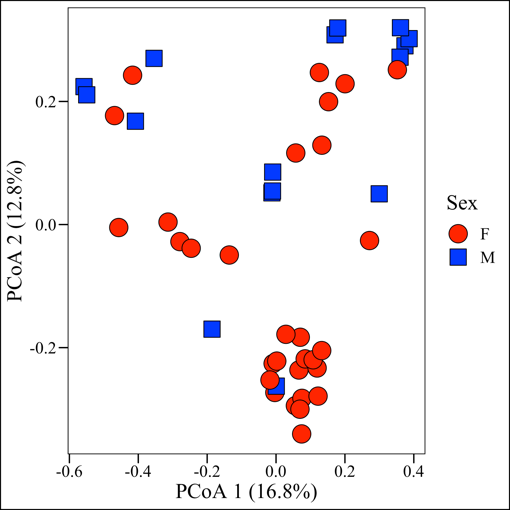

**d**

**e**

**f**

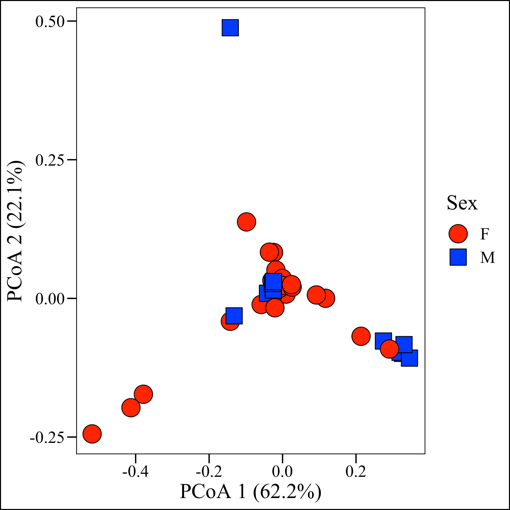

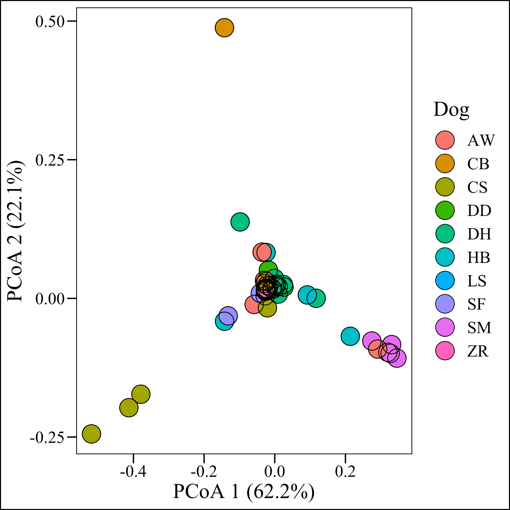

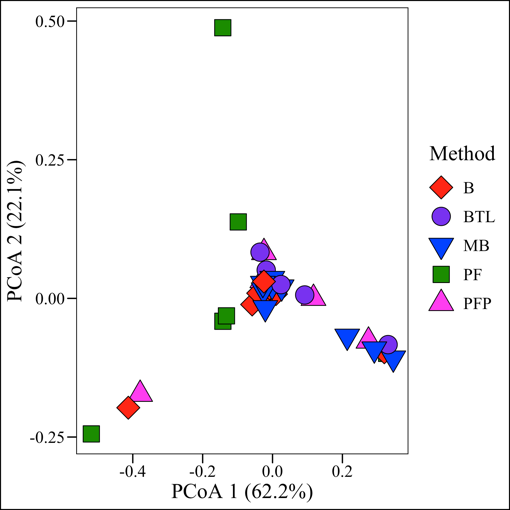

**Supplemental Figure 4 – Microbial Composition.** Unweighted (a, b, c) and Weighted UniFrac matrices (beta-diversity) (d, e, f) comparing microbial composition by (**a, d**) extraction method, (**b, e**) sex, (**c, f**) and dog. Microbial composition did not differ significantly by extraction method (Kruskal-Wallis: Unweighted UniFrac, *p* = 0.478; Weighted UniFrac, *p* = 0.524) but did differ significantly by sex (Kruskal-Wallis: Unweighted UniFrac, *p* = 0.005; Weighted UniFrac, *p* = 0.011) and dog (Kruskal-Wallis: Bray Curtis, *p* = < 0.001, Unweighted UniFrac, *p* = 0.005; Weighted UniFrac, *p* = 0.011)

**Figure S5 – Differentially Abundant Taxa by Sex**

**b**

**a**

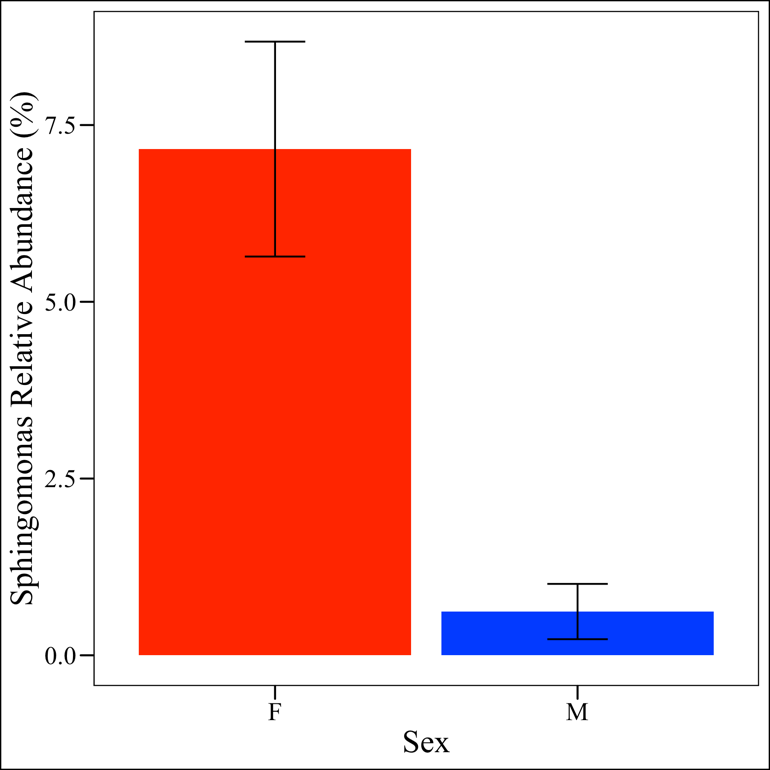

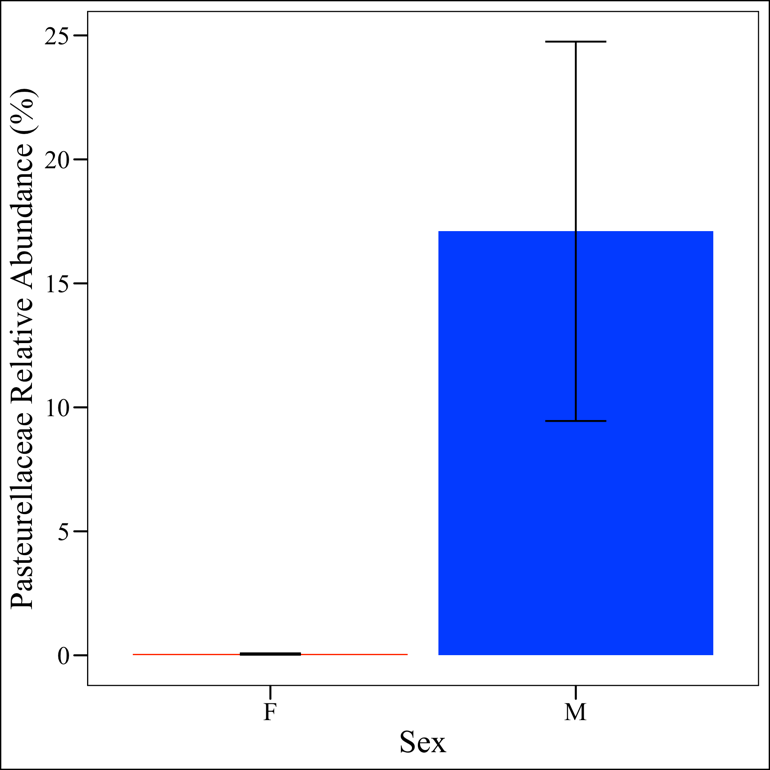

**Supplemental Figure 5 – Differentially Abundant Taxa by Sex**. (**a)** Females had significantly greater relative abundances of *Sphingomonas* (ANCOM, *W* = 573) while (**b)** males had significantly greater relative abundances of Pasteurellaceae bacterium canine oral taxon 272 (ANCOM, *W* = 591).

**Table S1 *-*** Dog Metadata

| **Dog ID** | **Sex** | **Spayed / Neutered?** | **Age (years)** | **Breed** |
| --- | --- | --- | --- | --- |
| AW | F | Y | 10 | Pit Bull / Boxer Mix |
| CB | M | Y | 2 | Golden Retriever / Border Collie |
| CS | F | Y | 1.5 | German Shepard / Belgian Malinois |
| DD | F | Y | 5 | Husky / Cattle Dog |
| DH | F | Y | 4 | Mastiff / St. Bernard Mix |
| HB | F | Y | 1.3 | Border Collie Mix |
| LS | F | N | 0.75 | Golden Retriever |
| SF | M | Y | 6 | Great Pyrenees |
| SM | M | N | 0.6 | Golden Retriever |
| ZR | M | Y | 1 | Pit Bull Mix |

**Table S2** *–* **Sample Concentrations and Reads.** Total and bacterial DNA concentrations and number of 16S reads in each sample. B = Bacteremia, BTL = Blood & Tissue with Lysozyme, MB = Magnetic Beads, PF = PowerFecal, PFP = PowerFecal Pro. * = Excluded from qPCR analysis as only one replicate amplified in qPCR. ** = Variation between replicates was greater than 3% but samples were included in analysis. *** Excluded from 16S analysis for having < 300 reads. Blue highlights = no bacterial DNA detected although total DNA was detected.

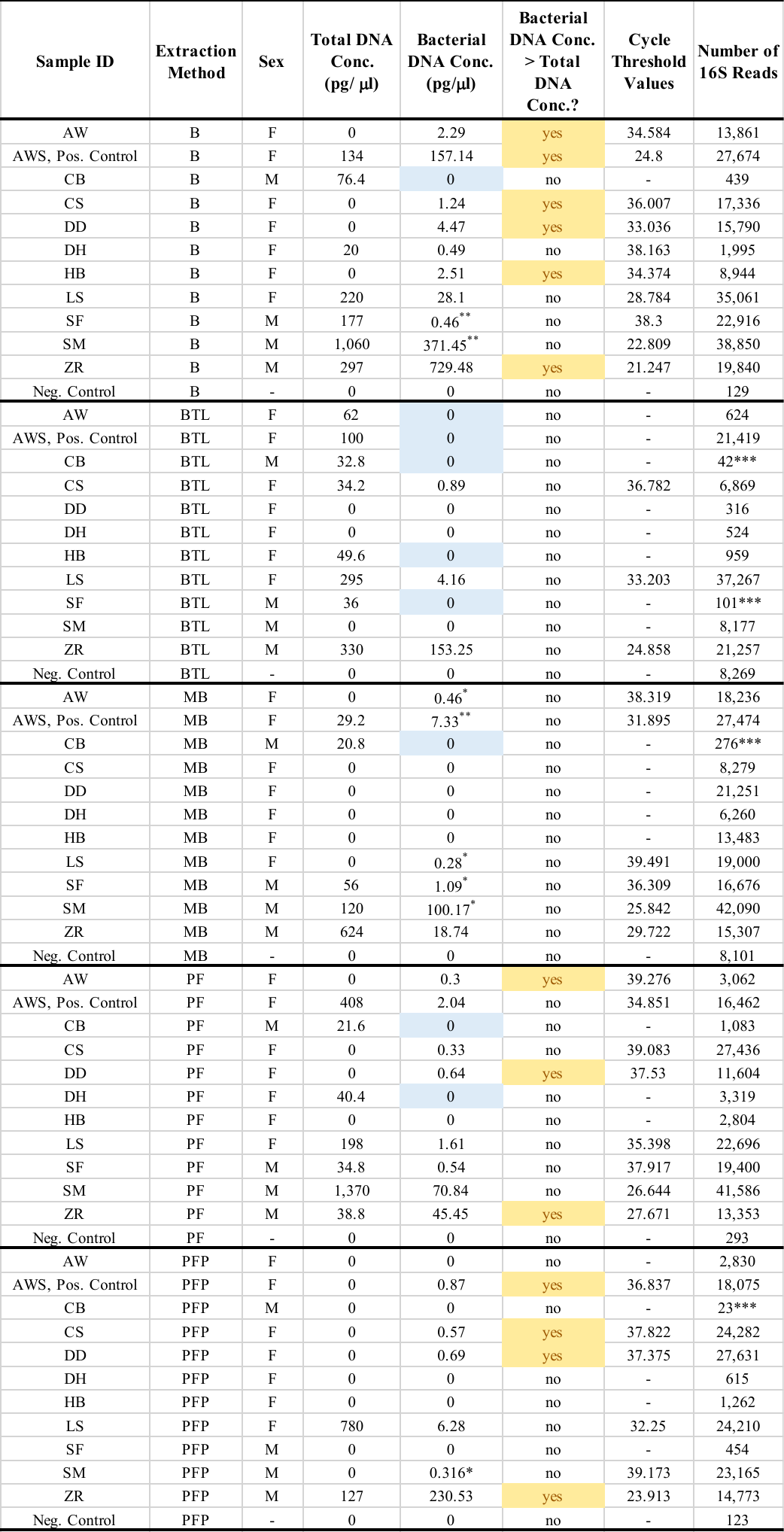

**Table S3 *–* Total DNA Concentration Pairwise Comparisons by Dog.** P-values based on Wilcoxon Rank Sum Tests for total DNA concentrations using 1000 permutations and False Discovery Rate corrections. There were no statistically significant pairwise comparisons.

|  | AW | AWS | CB | CS | CTL | DD | DH | HB | LS | SF | SM |
| --- | --- | --- | --- | --- | --- | --- | --- | --- | --- | --- | --- |
| AWS | 0.25 | - | - | - | - | - | - | - | - | - | - |
| CB | 0.29 | 0.48 | - | - | - | - | - | - | - | - | - |
| CS | 1.00 | 0.20 | 0.25 | - | - | - | - | - | - | - | - |
| CTL | 0.52 | 0.11 | 0.11 | 0.52 | - | - | - | - | - | - | - |
| DD | 0.52 | 0.11 | 0.11 | 0.52 | - | - | - | - | - | - | - |
| DH | 0.88 | 0.25 | 0.37 | 0.70 | 0.29 | 0.29 | - | - | - | - | - |
| HB | 1.00 | 0.25 | 0.29 | 1.00 | 0.52 | 0.52 | 0.88 | - | - | - | - |
| LS | 0.15 | 0.38 | 0.25 | 0.15 | 0.11 | 0.11 | 0.20 | 0.15 | - | - | - |
| SF | 0.29 | 0.98 | 0.48 | 0.15 | 0.11 | 0.11 | 0.29 | 0.25 | 0.29 | - | - |
| SM | 0.29 | 0.77 | 0.62 | 0.29 | 0.20 | 0.20 | 0.39 | 0.29 | 1.00 | 0.62 | - |
| ZR | 0.11 | 0.25 | 0.11 | 0.11 | 0.11 | 0.11 | 0.11 | 0.11 | 0.45 | 0.13 | 0.90 |

**Table S4 *–* Bacterial DNA concentration pairwise comparisons by dog.** P-values based on Wilcoxon Rank Sum Tests for total DNA concentrations using 1000 permutations and False Discovery Rate corrections. There were no statistically significant pairwise comparisons.

|  | AW | AWS | CB | CS | CTL | DD | DH | HB | LS | SF | SM |
| --- | --- | --- | --- | --- | --- | --- | --- | --- | --- | --- | --- |
| AWS | 0.341 | - | - | - | - | - | - | - | - | - | - |
| CB | 0.171 | 0.103 | - | - | - | - | - | - | - | - | - |
| CS | 0.610 | 0.405 | 0.103 | - | - | - | - | - | - | - | - |
| CTL | 0.171 | 0.103 | - | 0.103 | - | - | - | - | - | - | - |
| DD | 0.722 | 0.341 | 0.171 | 1.000 | 0.171 | - | - | - | - | - | - |
| DH | 0.497 | 0.139 | 0.530 | 0.171 | 0.530 | 0.293 | - | - | - | - | - |
| HB | 0.567 | 0.241 | 0.530 | 0.410 | 0.530 | 0.497 | 1.000 | - | - | - | - |
| LS | 0.119 | 0.778 | 0.086 | 0.094 | 0.086 | 0.171 | 0.094 | 0.107 | - | - | - |
| SF | 1.000 | 0.221 | 0.251 | 0.410 | 0.251 | 0.502 | 0.562 | 0.722 | 0.107 | - | - |
| SM | 0.497 | 0.722 | 0.171 | 0.556 | 0.171 | 0.497 | 0.302 | 0.302 | 0.717 | 0.497 | - |
| ZR | 0.086 | 0.164 | 0.086 | 0.086 | 0.086 | 0.086 | 0.086 | 0.086 | 0.109 | 0.103 | 0.824 |

**Table S5** – **Alpha-diversity pairwise comparisons by dog**. P-values resulting from pairwise Wilcoxon Rank Sum Tests for alpha diversity metrics using 1000 permutations

and False Discovery Rate corrections. *p < 0.05

| **Observed OTU p-values** | | | | | | | | | | | | |
| --- | --- | --- | --- | --- | --- | --- | --- | --- | --- | --- | --- | --- |
|  | AW | AWS | CB | CS | DD | DH | HB | | LS | | SF | SM |
| AWS | 0.073 | - | - | - | - | - | - | | - | | - | - |
| CB | 0.149 | 0.622 | - | - | - | - | - | | - | | - | - |
| CS | 0.415 | 0.044* | 0.149 | - | - | - | - | | - | | - | - |
| DD | 0.322 | 0.095 | 0.420 | 0.415 | - | - | - | | - | | - | - |
| DH | 0.092 | 0.877 | 0.877 | 0.044* | 0.133 | - | - | | - | | - | - |
| HB | 0.317 | 0.317 | 0.420 | 0.044* | 0.172 | 0.430 | - | | - | | - | - |
| LS | 0.044* | 0.044* | 0.149 | 0.044* | 0.133 | 0.044* | 0.044* | | - | | - | - |
| SF | 0.869 | 0.467 | 0.563 | 0.624 | 0.403 | 0.493 | 0.783 | | 0.092 | | - | - |
| SM | 0.322 | 0.088 | 0.274 | 0.044* | 0.268 | 0.285 | 1.000 | | 0.044* | | 0.922 | - |
| ZR | 0.044* | 0.149 | 0.149 | 0.044* | 0.044* | 0.149 | 0.095 | | 0.044* | | 0.285 | 0.044* |
| **Shannon p-values** | | | | | | | | | | | | |
|  | AW | AWS | CB | CS | DD | DH | HB | LS | | SF | | SM |
| AWS | 0.022* | - | - | - | - | - | - | - | | - | | - |
| CB | 0.146 | 0.146 | - | - | - | - | - | - | | - | | - |
| CS | 0.378 | 0.022* | 0.428 | - | - | - | - | - | | - | | - |
| DD | 0.889 | 0.022* | 0.428 | 0.378 | - | - | - | - | | - | | - |
| DH | 0.065 | 0.022* | 0.889 | 0.378 | 0.105 | - | - | - | | - | | - |
| HB | 0.313 | 0.022* | 0.428 | 1.000 | 0.146 | 0.591 | - | - | | - | | - |
| LS | 0.022* | 0.022* | 0.146 | 0.022* | 0.105 | 0.022* | 0.022* | - | | - | | - |
| SF | 0.038* | 0.038* | 0.587 | 0.165 | 0.065 | 0.276 | 0.116 | 0.038 | | - | | - |
| SM | 0.022* | 0.022* | 0.428 | 1.000 | 0.146 | 0.378 | 0.378 | 0.022* | | 0.378 | | - |
| ZR | 0.022* | 0.065 | 0.146 | 0.022* | 0.022* | 0.022* | 0.022* | 0.022* | | 0.065 | | 0.022* |
| **Simpson p-values** | | | | | | | | | | | | |
|  | AW | AWS | CB | CS | DD | DH | HB | | LS | | SF | SM |
| AWS | 0.026* | - | - | - | - | - | - | | - | | - | - |
| CB | 0.150 | 0.150 | - | - | - | - | - | | - | | - | - |
| CS | 0.105 | 0.026* | 0.889 | - | - | - | - | | - | | - | - |
| DD | 0.387 | 0.026* | 0.437 | 0.387 | - | - | - | | - | | - | - |
| DH | 0.038* | 0.026* | 0.437 | 0.615 | 0.105 | - | - | | - | | - | - |
| HB | 0.105 | 0.026* | 0.437 | 0.730 | 1.000 | 0.387 | - | | - | | - | - |
| LS | 0.730 | 0.026* | 0.150 | 0.150 | 0.313 | 0.026* | 0.026* | | - | | - | - |
| - | 0.038* | 0.038* | 0.198 | 0.116 | 0.038* | 0.170 | 0.067 | | 0.038 | | - | - |
| SM | 0.038* | 0.026* | 0.437 | 1.000 | 0.730 | 0.387 | 0.313 | | 0.026* | | 0.067 | - |
| ZR | 0.026* | 0.067 | 0.150 | 0.026* | 0.026* | 0.026* | 0.026* | | 0.026* | | 0.387 | 0.026* |

**Table S6** – **Beta-diversity pairwise comparisons by dog**. P-values based on PERMANOVA pairwise comparisons using 1000 permutations False Discovery Rate corrections. * = p < 0.05

| **Bray Curtis p-values** | | | | | | | | | | |
| --- | --- | --- | --- | --- | --- | --- | --- | --- | --- | --- |
|  | DH | LS | HB | SM | AW | AWS | ZR | SF | CS | DD |
| CB | 0.339 | 0.0583 | 0.109 | 0.0583 | 0.119 | 0.0561 | 0.0604 | 0.138 | 0.05 | 0.0525 |
| DD | 0.0366^*^ | 0.0206^*^ | 0.0206^*^ | 0.0220^*^ | 0.0206^*^ | 0.0220^*^ | 0.0206^*^ | 0.0316^*^ | 0.0206^*^ |  |
| CS | 0.0206^*^ | 0.0206^*^ | 0.0206^*^ | 0.0206^*^ | 0.0206^*^ | 0.0206^*^ | 0.0206^*^ | 0.0206^*^ |  |  |
| SF | 0.0362^*^ | 0.0206^*^ | 0.0206^*^ | 0.0206^*^ | 0.0220^*^ | 0.0206^*^ | 0.0206^*^ |  |  |  |
| ZR | 0.0206^*^ | 0.0206^*^ | 0.0206^*^ | 0.0206^*^ | 0.0206^*^ | 0.0206^*^ |  |  |  |  |
| AWS | 0.0206^*^ | 0.0206^*^ | 0.0206^*^ | 0.0206^*^ | 0.0206^*^ |  |  |  |  |  |
| AW | 0.0473^*^ | 0.0206^*^ | 0.263 | 0.0238^*^ |  |  |  |  |  |  |
| SM | 0.0229^*^ | 0.0206^*^ | 0.0282^*^ |  |  |  |  |  |  |  |
| HB | 0.0583 | 0.0206^*^ |  |  |  |  |  |  |  |  |
| LS | 0.0206^*^ |  |  |  |  |  |  |  |  |  |
| **Unweighted UniFrac p-values** | | | | | | | | | | |
|  | DH | LS | HB | SM | AW | AWS | ZR | SF | CS | DD |
| CB | 0.538 | 0.0792 | 0.468 | 0.0792 | 0.543 | 0.0792 | 0.540 | 0.468 | 0.0792 | 0.0739 |
| DD | 0.09 | 0.043^*^ | 0.142 | 0.0325^*^ | 0.204 | 0.09 | 0.0305^*^ | 0.251 | 0.0305^*^ |  |
| CS | 0.0440^*^ | 0.0318^*^ | 0.0305^*^ | 0.0305^*^ | 0.0687 | 0.0325^*^ | 0.0305^*^ | 0.0305^*^ |  |  |
| SF | 0.426 | 0.0305^*^ | 0.540 | 0.0305^*^ | 0.543 | 0.0687 | 0.251 |  |  |  |
| ZR | 0.503 | 0.0305^*^ | 0.220 | 0.0305^*^ | 0.126 | 0.0613 |  |  |  |  |
| AWS | 0.312 | 0.0305^*^ | 0.0792 | 0.0305^*^ | 0.0305^*^ |  |  |  |  |  |
| AW | 0.265 | 0.0305^*^ | 0.540 | 0.0305^*^ |  |  |  |  |  |  |
| SM | 0.0305^*^ | 0.0325^*^ | 0.0440^*^ |  |  |  |  |  |  |  |
| HB | 0.704 | 0.0305^*^ |  |  |  |  |  |  |  |  |
| LS | 0.0305^*^ |  |  |  |  |  |  |  |  |  |
| **Weighted UniFrac p-values** | | | | | | | | | | |
|  | DH | LS | HB | SM | AW | AWS | ZR | SF | CS | DD |
| CB | 0.248 | 0.119 | 0.289 | 0.107 | 0.246 | 0.0989 | 0.427 | 0.306 | 0.246 | 0.117 |
| DD | 0.749 | 0.333 | 0.573 | 0.0353^*^ | 0.573 | 0.0353^*^ | 0.0388^*^ | 0.0353^*^ | 0.0989 |  |
| CS | 0.156 | 0.220 | 0.156 | 0.0366^*^ | 0.218 | 0.117 | 0.256 | 0.246 |  |  |
| SF | 0.449 | 0.0353^*^ | 0.578 | 0.0353^*^ | 0.573 | 0.0353^*^ | 0.0353^*^ |  |  |  |
| ZR | 0.835 | 0.0353^*^ | 0.934 | 0.0353^*^ | 0.905 | 0.0353^*^ |  |  |  |  |
| AWS | 0.385 | 0.0353^*^ | 0.548 | 0.0353^*^ | 0.222 |  |  |  |  |  |
| AW | 0.857 | 0.726 | 0.857 | 0.0672 |  |  |  |  |  |  |
| SM | 0.0353^*^ | 0.0378^*^ | 0.0353^*^ |  |  |  |  |  |  |  |
| HB | 0.950 | 0.573 |  |  |  |  |  |  |  |  |
| LS | 0.726 |  |  |  |  |  |  |  |  |  |

**Table S7 – Differentially abundant taxa by dog**. Thirty-seven taxa at the L7 level were differentially abundant by dog.

| **Taxa ID** | **W** |
| --- | --- |
| D_1 Firmicutes;D_2 Bacilli;D_3 Lactobacillales;D_4 Enterococcaceae;D_5 Enterococcus;__ | 622 |
| D_1 Firmicutes;D_2 Bacilli;D_3 Lactobacillales;D_4 Streptococcaceae;D_5 Streptococcus;D_6 Streptococcus canis | 622 |
| D_1 Firmicutes;D_2 Bacilli;D_3 Lactobacillales;D_4 Streptococcaceae;D_5 Streptococcus;D_6 Streptococcus halichoeri | 622 |
| D_1 Proteobacteria;D_2 Gammaproteobacteria;D_3 Pasteurellales;D_4 Pasteurellaceae;D_5 Haemophilus;D_6 Pasteurellaceae bacterium canine oral taxon 272 | 622 |
| D_1 Proteobacteria;D_2 Gammaproteobacteria;D_3 Pasteurellales;D_4 Pasteurellaceae;D_5 Haemophilus;__ | 622 |
| D_1 Proteobacteria;D_2 Gammaproteobacteria;D_3 Pseudomonadales;D_4 Moraxellaceae;D_5 Psychrobacter;D_6 Psychrobacter alimentarius | 622 |
| D_1 Firmicutes;D_2 Bacilli;D_3 Bacillales;D_4 Family XII;D_5 Exiguobacterium;__ | 620 |
| D_1 Proteobacteria;D_2 Gammaproteobacteria;D_3 Pseudomonadales;D_4 Pseudomonadaceae;D_5 Pseudomonas;__ | 618 |
| D_1 Tenericutes;D_2 Mollicutes;D_3 Mycoplasmatales;D_4 Mycoplasmataceae;D_5 Mycoplasma;D_6 Mycoplasma cynos C142 | 613 |
| D_1 Fusobacteria;D_2 Fusobacteriia;D_3 Fusobacteriales;D_4 Leptotrichiaceae;D_5 Oceanivirga;D_6 uncultured bacterium | 612 |
| D_1 Tenericutes;D_2 Mollicutes;D_3 Mycoplasmatales;D_4 Mycoplasmataceae;D_5 Mycoplasma;__ | 612 |
| D_1 Tenericutes;D_2 Mollicutes;D_3 Mycoplasmatales;D_4 Mycoplasmataceae;D_5 Ureaplasma;D_6 Ureaplasma canigenitalium | 612 |
| D_1 Bacteroidetes;D_2 Bacteroidia;D_3 Sphingobacteriales;D_4 Sphingobacteriaceae;D_5 Sphingobacterium;__ | 611 |
| D_1 Fusobacteria;D_2 Fusobacteriia;D_3 Fusobacteriales;D_4 Fusobacteriaceae;D_5 Fusobacterium;D_6 Fusobacterium sp. canine oral taxon 439 | 611 |
| D_1 Tenericutes;D_2 Mollicutes;D_3 Acholeplasmatales;D_4 Acholeplasmataceae;D_5 Acholeplasma;D_6 Mycoplasma feliminutum | 610 |
| D_1 Bacteroidetes;D_2 Bacteroidia;D_3 Bacteroidales;D_4 Prevotellaceae;D_5 Alloprevotella;D_6 Prevotella sp. canine oral taxon 226 | 609 |
| D_1 Firmicutes;D_2 Clostridia;D_3 Clostridiales;D_4 Family XI;D_5 Parvimonas;__ | 607 |
| D_1 Firmicutes;D_2 Clostridia;D_3 Clostridiales;D_4 Lachnospiraceae;D_5 Acetitomaculum;D_6 uncultured bacterium | 607 |
| D_1 Firmicutes;D_2 Clostridia;D_3 Clostridiales;D_4 Peptococcaceae;D_5 Peptococcus;__ | 607 |
| D_1 Bacteroidetes;D_2 Bacteroidia;D_3 Bacteroidales;D_4 Tannerellaceae;D_5 Parabacteroides;__ | 606 |
| D_1 Firmicutes;D_2 Erysipelotrichia;D_3 Erysipelotrichales;D_4 Erysipelotrichaceae;D_5 Solobacterium;D_6 uncultured Erysipelotrichaceae bacterium | 606 |
| D_1 Firmicutes;D_2 Bacilli;D_3 Bacillales;D_4 Family XI;D_5 Gemella;__ | 605 |
| D_1 Firmicutes;D_2 Clostridia;D_3 Clostridiales;D_4 Peptostreptococcaceae;D_5 Peptostreptococcus;__ | 605 |
| D_1 Fusobacteria;D_2 Fusobacteriia;D_3 Fusobacteriales;D_4 Leptotrichiaceae;D_5 Streptobacillus;__ | 602 |
| D_1 Proteobacteria;D_2 Gammaproteobacteria;D_3 Enterobacteriales;D_4 Enterobacteriaceae;D_5 Pantoea;__ | 599 |
| D_1 Actinobacteria;D_2 Actinobacteria;D_3 Actinomycetales;D_4 Actinomycetaceae;D_5 Actinomyces;D_6 Actinomyces sp. canine oral taxon 374 | 597 |
| D_1 Proteobacteria;D_2 Alphaproteobacteria;D_3 Sphingomonadales;D_4 Sphingomonadaceae;D_5 Novosphingobium;__ | 596 |
| D_1 Bacteroidetes;D_2 Bacteroidia;D_3 Flavobacteriales;D_4 Flavobacteriaceae;D_5 Flavobacterium;D_6 Flavobacterium ummariense | 588 |
| D_1 Proteobacteria;D_2 Gammaproteobacteria;D_3 Enterobacteriales;D_4 Enterobacteriaceae;__;__ | 583 |
| D_1 Bacteroidetes;D_2 Bacteroidia;D_3 Sphingobacteriales;D_4 Sphingobacteriaceae;D_5 Pedobacter;__ | 582 |
| D_1 Bacteroidetes;D_2 Bacteroidia;D_3 Flavobacteriales;D_4 Weeksellaceae;D_5 Chryseobacterium;__ | 577 |
| D_1 Proteobacteria;D_2 Gammaproteobacteria;D_3 Enterobacteriales;D_4 Enterobacteriaceae;D_5 Escherichia-Shigella;__ | 573 |
| D_1 Proteobacteria;D_2 Alphaproteobacteria;D_3 Rhizobiales;D_4 Devosiaceae;D_5 Devosia;__ | 572 |
| D_1 Proteobacteria;D_2 Alphaproteobacteria;D_3 Rhizobiales;D_4 Rhizobiaceae;D_5 Allorhizobium-Neorhizobium-Pararhizobium-Rhizobium;__ | 567 |
| D_1 Proteobacteria;D_2 Gammaproteobacteria;D_3 Cardiobacteriales;D_4 Cardiobacteriaceae;D_5 Suttonella;D_6 Rappaport israeli | 565 |
| D_1 Bacteroidetes;D_2 Bacteroidia;D_3 Flavobacteriales;D_4 Flavobacteriaceae;D_5 Flavobacterium;__ | 563 |
| D_1 Proteobacteria;D_2 Gammaproteobacteria;D_3 Pseudomonadales;D_4 Moraxellaceae;D_5 Psychrobacter;__ | 561 |
